# Ciliary Genes *arl13b*, *ahi1* and *cc2d2a* Differentially Modify Expression of Visual Acuity Phenotypes but do not Enhance Retinal Degeneration due to Mutation of *cep*290 in Zebrafish

**DOI:** 10.1101/569822

**Authors:** Emma M. Lessieur, Ping Song, Gabrielle C. Nivar, Ellen M. Piccillo, Joseph Fogerty, Richard Rozic, Brian D. Perkins

## Abstract

Mutations in the gene Centrosomal Protein 290 kDa (*CEP290*) result in multiple ciliopathies ranging from the neonatal lethal disorder Meckel-Gruber Syndrome to multi-systemic disorders such as Joubert Syndrome and Bardet-Biedl Syndrome to nonsyndromic diseases like Leber Congenital Amaurosis (LCA) and retinitis pigmentosa. Results from model organisms and human genetics studies, have suggest that mutations in genes encoding protein components of the transition zone (TZ) and other cilia-associated proteins can function as genetic modifiers and be a source for *CEP290* pleiotropy. We investigated the zebrafish *cep290*^*fh297/fh297*^ mutant, which encodes a nonsense mutation (p.Q1217*). This mutant is viable as adults, exhibits scoliosis, and undergoes a slow, progressive cone degeneration. The *cep290*^*fh297/fh297*^ mutants showed partial mislocalization of the transmembrane protein rhodopsin but not of the prenylated proteins rhodopsin kinase (GRK1) or the rod transducin subunit GNB1. Surprisingly, photoreceptor degeneration did not trigger proliferation of Müller glia, but proliferation of rod progenitors in the outer nuclear layer was significantly increased. To determine if heterozygous mutations in other cilia genes could exacerbate retinal degeneration, we bred *cep290*^*fh297/fh297*^ mutants to *arl13b, ahi1,* and *cc2d2a* mutant zebrafish lines. While *cep290*^*fh297/fh297*^ mutants lacking a single allele of these genes did not exhibit accelerated photoreceptor degeneration, loss of one alleles of *arl13b* or *ahi1* reduced visual performance in optokinetic response assays at 5 days post fertilization. Our results indicate that the *cep290*^*fh297/fh297*^ mutant is a useful model to study the role of genetic modifiers on photoreceptor degeneration in zebrafish and to explore how progressive photoreceptor degeneration influences regeneration in adult zebrafish.

**Nonstandard abbreviations:**

BBS: Bardet-Biedl Syndrome
COS: cone outer segments
Dpf: Days post fertilization
GNB1: rod transducin β subunit
GRK1: rhodopsin kinase
JTBS: Joubert Syndrome
LCA: Leber Congenital Amaurosis
MKS: Meckel Syndrome
NPHP: nephronophthisis
OKR: optokinetic response
PNA: peanut agglutinin lectin
ROS: rod outer segments
RP2: Retinitis Pigmentosa 2

## INTRODUCTION

Ciliopathies refer to a group of recessive disorders stemming from defects in the biogenesis, structure or function of cilia [1]. These disorders exhibit both clinical and genetic heterogeneity [2], with mutations in dozens of genes resulting in a spectrum of diseases sharing overlapping symptoms. Clinically, ciliopathies can manifest as non-syndromic disorders, such as in Leber Congenital Amaurosis (LCA; OMIM: 204000), which is an inherited form of childhood retinal dystrophy, to more pleiotropic diseases, such as Joubert Syndrome (JBTS; OMIM 213300), Meckel Syndrome (MKS; OMIM 249000), Bardet-Biedl Syndrome (BBS; OMIM 209900), and Nephronophthisis (NPHP; OMIM 256100), each of which impact unique combinations of organ systems [3].

Mutations in the gene for Centrosomal Protein 290 (*CEP290*) cause JBTS, MKS, BBS, and NPHP [3], but also account for 15-25% of cases of isolated blindness in LCA with no associated systemic disease [4, 5]. Most *CEP290* lesions in humans are stop codons that result from frameshift and nonsense mutations occurring throughout the gene, whereas pathogenic missense mutations are rare [6]. Despite the identification of more than 130 mutations in human *CEP290*, efforts to establish obvious genotype-phenotype correlations have been unsuccessful [6]. *CEP290* mutations are strongly associated with retinal dystrophy and photoreceptor degeneration is one of the most common symptoms of ciliopathies [7]. The most frequent *CEP290* mutation in LCA is a deep intronic mutation (c.2991+1655 A>G) that activates a cryptic splice site and creates a stop codon, resulting in early termination of the protein [4]. In a recent study of LCA patients with *CEP290* mutations, visual acuity varied considerably although most patients had significant vision loss and undetectable electroretinograms (ERGs) regardless of genotype [8]. In spite of the severe loss of vision, multiple studies have reported that cone photoreceptors persist within the central retina of *CEP290*-LCA patients [8-11], suggesting that a window of opportunity may exist for therapeutic intervention.

In humans, *CEP290* encodes a 2479 amino acid protein (∼290 kDa) that can localize to the basal body [12] and/or the transition zone (TZ) of cilia [13, 14] in a tissue-specific manner. The TZ refers to the most proximal region of the ciliary axoneme, immediately distal to the basal body. The connecting cilium in vertebrate photoreceptors is homologous to the TZ of a prototypic primary cilium [15]. The TZ is believed to function as a ciliary gate that regulates protein entry and exit to the cilium. Defects in the ciliary gate may result in abnormal accumulation of non-outer segment proteins within the outer segment [16] and/or disrupt normal protein delivery to the outer segment. Work from *C. elegans* have proposed roles for Cep290 ranging from cell adhesion to TZ assembly [13, 17, 18], but *in vivo* studies in vertebrate models have not fully elucidated a role for Cep290 or explained the variability in photoreceptor phenotypes [14, 19-21]. Abyssinian cats exhibit a high degree of inherited retinal degeneration due to the *rdAc* allele in the feline *Cep290* gene and this *rdAc* allele can also be found at elevated frequencies in several other cat breeds [22, 23]. In rodents, the *rd16* allele reflects an in-frame deletion of *Cep290* that leads to rapid photoreceptor degeneration [20]. Although a targeted gene knockout of *Cep290* causes embryonic lethality in mice [14], truncating nonsense mutations in humans can often result in attenuated pathologies that range from multisystem dysfunction to mild retinal disease. Indeed, LCA patients with two truncating *CEP290* mutations can sometimes maintain photoreceptor architecture and retain limited visual acuity [8, 9, 24]. These unexpectedly mild phenotypes have recently been attributed to basal exon skipping and nonsense-mediated alternative splicing of the *CEP290* mRNA [24, 25]. Nevertheless, patients with identical genotypes can still exhibit very different retinal phenotypes [9].

One hypothesis to explain the variable phenotypic expression is the effects of mutations in second-site modifiers [26-29]. Genetic [26-28] and biochemical studies [18, 30] in *C. elegans* and cultured mammalian cells have identified two molecular complexes within the TZ, termed the NPHP and MKS modules. The proteins Cc2d2a, Ahi1, Mks1, and at least 5 other proteins form the MKS module [28], while the NPHP module consists of Nphp1 and Nphp4 [31], although proteomic studies suggest additional factors likely exist [30]. Homozygous mutations in genes from both an MKS and NPHP module severely disrupt cilia formation [27, 28, 31, 32]. These genetic interactions, however, reflect phenotypes resulting from double homozygous mutants. In humans, the frequency of pathogenic alleles in cilia genes is low and a more realistic scenario is that heterozygous mutant alleles in one cilia gene may influence phenotypic expression resulting from homozygous mutations in *CEP290*. Heterozygous missense alleles of the *AHI1* gene were identified in LCA patients with severe neurological involvement, suggesting that alleles of *AHI1* may influence phenotypic expression [33]. In zebrafish, morpholino suppression of *cep290* resulted in a genetic interaction with *cc2d2a* and synergistically enhanced kidney cyst phenotypes [12]. Finally, the small GTPase Arl13b localizes to cilia and is essential for photoreceptor survival [34, 35]. The *C. elegans* homolog *arl-13* genetically interacts with *nphp-2* to regulate ciliogenesis and Arl13b was reported to regulate cilia length [36]. Genetic interactions between *ARL13B* and other ciliary components have not been investigated; however, protein complexes containing Cep290 show similar localization patterns with Arl13b to the basal body and TZ domains.

In this study, we evaluated the zebrafish *cep290*^*fh297/fh297*^ mutant in an effort to test *ahi1, cc2d2a,* and *arl13b* as potential genetic modifiers of retinal degeneration. We report that the *cep290*^*fh297/fh297*^ mutant shows progressive and predominant cone degeneration. We found that the phenotype observed in these mutants was not the consequence of nonsense-associated alternative-splicing, a phenomenon hypothesized to explain phenotypic variation in humans [37]. We report that heterozygous mutations in *ahi1* and *arl13b*, were associated with decreased visual acuity, whereas the absence of one allele of *cc2d2a* had no effect on visual acuity. Retinal degeneration in the *cep290*^*fh297/fh297*^ mutant was not exacerbated by heterozygosity of any of these genes. Furthermore, the mild phenotypes observed in *cep290*^*fh297/fh297*^ mutants was not due to retinal regeneration These data demonstrate a role for Cep290 in cone survival in zebrafish and provide a foundation for future analysis of potential modifier genes of *cep290*- associated retinal degeneration.

## MATERIAL AND METHODS

### Zebrafish husbandry

Adult zebrafish were maintained and raised on an Aquatic Habitats recirculating water system (Pentair; Apopka, FL, USA) in a 14:10 hr light-dark cycle. The Cleveland Clinic Institutional Animal Care and Use Committee (IACUC) approved all experimental procedures (Protocol number: 2018-1980). The *cep290*^*fh297/fh297*^ mutant was identified by the zebrafish TILLING consortium and was a gift of Dr. Cecilia Moens (Fred Hutchinson Cancer Center, Seattle, WA. USA).

### Sequencing

Using sequence data from Ensembl (http://useast.ensembl.org/Danio_rerio/Info/Index; transcript: *cep290-202*), primers were designed to span exon 29 (5′-GTCTGATGAAAAGGCCCTGA-3’ and 5’-CCTCCAAGCCTTTCAGCTTT-3’) for the *cep290*^*fh297*^ allele. Samples were sequenced at the Genomics Core of the Cleveland Clinic Lerner Research Institute using the high-throughput, 96-capillary *ABI 3730*x/DNA Analyzer.

### Genotyping

c*DNA extraction:* Tail-clips from embryos or adults were placed in 0.5 ml individual tubes and 25 μL (embryos) or 50 μL (adults) of lysis buffer (50mM Tris pH 8.5, 5mM EDTA, 100mM NaCl, 0.4% SDS, 100 μg/mL proteinase K) was added to each tube and then incubated at 60 °C for 4 hrs (embryos) to overnight (adults). Samples were then diluted 1:10 in nuclease-free water and heat inactivated at 95 °C for 5 min.

*PCR and High Resolution Melt Analysis (HRMA)*: HRMA primers targeting *cep290* exon 29 for the *cep290*^*fh297*^ allele (5′ - ACAAACACACGTCTGCAGAAACTGGACGCG – 3’ and 5’ -CTGCTGTTGCTCATCCAG TT – 3’) were designed flanking the point mutation. PCR products were 95 bp. High-resolution melt curve analysis was performed using Bio-Rad Precision Melt reagent in 8μl reactions with a CFX96 Touch™ Real-Time PCR Detection System (Bio-Rad, Hercules, CA, USA) at standard cycling conditions. Melt curves were analyzed using the Precision Melt Analysis Software version 1.2 (Bio-Rad, Hercules, CA, USA).

### Micro-computed tomography (μCT)

Adult zebrafish were euthanized and fixed in 4% paraformaldehyde (in 1X PBS) overnight at 4 °C. Specimens were washed in 1X PBS and immersed in 70% ethanol in a 15 mL conical tube. Samples were scanned with an Explore Locus RS (GE Medical Systems, London, Ontario, Canada) at 45 μm. Images were analyzed and reconstructed using MicroView software version 2.5.0-3768 (Parallax Innovations Inc.; Ilderton, Ontario, Canada).

### Light and electron microscopy

Light-adapted larvae were bisected through the swim bladder, and heads were prepared for transmission electron microscopy. Tails were used for genomic DNA extraction and genotyping as described above. For adult animals, enucleations were performed at the designated time points and samples prepared for transmission electron microscopy. Briefly, the eyes were enucleated from light-adapted animals and the anterior segment was dissected away in primary fixative (0.08M cacodylate buffer containing 2% paraformaldehyde and 2% glutaraldehyde). The tissue was fixed for 1 hr at room temperature in primary fixative and then washed with cacodylate buffer and post-fixed in 1% osmium tetroxide for 1 hr at 4 °C. Samples were washed again and then dehydrated in a graded methanol series before embedding them in Embed-812/DER736 (Electron Microscopy Sciences; Hatfield, PA, USA), using acetonitrile as a transition solvent. Semi-thin sections were made with a Leica EM UC7 ultramicrotome (Leica Microsystems; GmbH Vienna, Austria), stained with Toluidine Blue, and imaged with a Zeiss Axio Imager.Z2 (Carl Zeiss Microscopy, Thornwood, NY, USA). Ultrathin sections were stained with uranyl acetate and lead citrate following standard procedures, and electron microscopy was performed on a Tecnai G2 Spirit BioTWIN 20-120 kV digital electron microscope (FEI Company; Hillsboro, OR, USA). Micrographs were acquired with a Gatan image filter and an Orius 832 CCD Camera (Gatan, Inc.; Pleasanton, CA, USA).

### Optokinetic Response (OKR)

OKR measurements on 5-6 dpf larvae were conducted between 12-6 pm using the VisioTracker system (VisioTracker 302060 Series, TSE Systems, GmbH Bad Homburg, Germany). Contrast sensitivity was assessed as described previously [38, 39]. For the spatial frequency response function [39, 40], the contrast was held constant at 70% and we tested stimuli of 0.02, 0.04, 0.06, 0.08, 0.12, and 0.16 cycles/degree by first increasing and then decreasing the frequency. Each spatial frequency stimulus was presented for 3 seconds before reversing direction for another 3 seconds to minimize saccade frequency. All OKR stimuli were presented with a constant angular velocity of 7.5 degrees per second. The genotypes of individual larvae were confirmed following OKR tests.

### Immunohistochemistry and fluorescence imaging

Larvae were fixed for 2 hrs. at 4 °C. Adult eyes were fixed at the designated time points. Fixation protocols varied depending on the primary antibodies being used. For Zpr-1, Zpr3, GRK1, and GNB1, samples were fixed in 4% paraformaldehyde in 0.8X PBS at 4 °C overnight. For peanut agglutinin (PNA) and acetylated tubulin staining, heads were fixed in 4% paraformaldehyde in 0.8X PBS at 4 °C for a maximum of 2 hrs. All samples were cryoprotected in 30% sucrose overnight. Cryosections (10 μM) were cut and dried at room temperature overnight. Blocking solution (1% BSA, 5% normal goat serum, 0.2% Triton-X-100, 0.1% Tween-20 in 1X PBS) was applied for 2 hrs in a dark, humidified chamber. Primary antibodies were diluted in blocking solution as follow: Zpr-1 and Zpr-3 (1:200; Zebrafish International Resource Center, University of Oregon, Eugene, OR, USA), GNB1 (1:100; Abgent AP5036a), GRK1 (1:50; Abclonal A6497), and acetylated-α-tubulin (1:5000; Sigma 6-11-B1). Conjugated secondary antibodies were purchased from Invitrogen Life Technologies (Carlsbad, CA, USA) and used at 1:500 dilutions and 4,6-diamidino-2-phenylendole (DAPI; 1:1000) was used to label nuclei. Optical sections were obtained with a Zeiss Axio Imager.Z2 fluorescent microscope fitted with the Apotome.2 for structured illumination (Carl Zeiss Microscopy, Thornhill, NY. USA). ImageJ was used to create image panels. Figures were assembled in Adobe Photoshop. To quantify cone outer segment density, the number of PNA-positive outer segments was determined from images of transverse sections of dorsal retinas. The distance of retina measured in each section was determined using ImageJ and density was calculated as the number of cone outer segments per 50 microns of retina. Each data point represented one section from a distinct retina. To quantify rhodopsin mislocalization, ImageJ was used to measure the integrated fluorescence density (IFD) across a region of interest (ROI) of defined area that was placed in the rod outer segments (ROS; proper localization), inner segment/outer nuclear layer (IS/ONL; mislocalized) or the inner nuclear layer (INL; background fluorescence). Corrected fluorescence intensities were calculated by subtracting the background fluorescence. The total rhodopsin fluorescence was calculated as the sum of the IFD from the ROS and IS/ONL. The percentage of mislocalized rhodopsin was calculated as the IFD from the IS/ONL (numerator) per total rhodopsin (denominator).

### RT-PCR

For traditional RT-PCR, total RNA was extracted from pooled wild-type larvae at 5 dpf for positive control (tunicamycin), and from 4 isolated retinas from wild-type and *cep290*^*fh297/fh297*^ mutants at the designated time points using TRIzol according to standard protocols. cDNA was reverse transcribed using SuperScript II reverse transcriptase and random hexamers according to the manufacturer’s instructions (ThermoFisher Scientific, Waltham, MA. USA). RT-PCR was performed according to standard protocols and cycling conditions.

*Exon skipping:* cDNA from wild-type and *cep290*^*fh297/fh297*^ mutants was obtained as described above. Primers were designed to encompass the mutated exon as follow: primers targeting exons 27-32 for *cep290*^*fh297/fh297*^ (5’ – AGAATCACTGAACTGGAGAAAACAG – 3’ and TTCCTTTTCTTTTAGCTTCTCTTCC – 3’) with products sizes – 1040 bp when mutated exon is included and 593 when mutated exon is skipped.

*Droplet digital PCR:* RNA was isolated from whole eyes of 6-month old *cep290* mutant and sibling control animals (n = 9 per group) with Trizol (ThermoFisher 15596026). Reverse transcription using 1 μg RNA was performed with the iScript cDNA Synthesis kit (Bio-Rad 1708891). Intron-spanning primers and probes for *cep290* and *ef1a1l1* were designed with Primer3Plus (http://primer3plus.com). Sequences are as follows: cep290F – ACACCGTCATCCAGCTGAAG; cep290R – CTGGCAAGACCTTCGTCAGT; cep290probe(FAM) – ACGTCCCTGTGGAAGCGACC; ef1a1l1F – CGTCTGCCACTTCAGGATGT; ef1a1l1R – CCCAGCCTTCAGAGTTCCAG; ef1a1l1probe(HEX) – ACTGTGCCTGTGGGACGTGT. Multiplexed PCR reactions using 100 ng cDNA were prepared with the ddPCR supermix for probes (No dUTP, Bio-Rad 1863024) and fractionated into >20,000 droplets using the Bio-Rad QX200 droplet generator with droplet generation oil for probes (Bio-Rad 1863005). PCR cycling was performed using a 60 degree C annealing temperature, and droplet signal was detected with a QX200 droplet reader (Bio-Rad). Target copy number was determined with QuantaSoft Analysis Pro software (Bio-Rad) after droplets were manually thresholded relative to no-template control reactions.

### Statistics

Graphs were generated using Prism6 (GraphPad Software; San Diego, USA). Statistical analyses were performed using a one-way ANOVA with a Multiple Comparisons test and Tukey’s correction or 2-way ANOVA with a Multiple Comparisons test and Sidek corrections. For all tests, P-values less than 0.05 were considered significant.

## RESULTS

### Identification of a nonsense mutation in zebrafish *cep290* gene

In zebrafish, the primary *cep290* transcript (RefSeq: NM_001168267) encodes a 2439 amino acid protein. The *cep290*^*fh297*^ allele was identified by the Zebrafish TILLING Consortium. This C>T transition mutation results in a stop codon (p.Gln1217X) downstream of the Cc2d2a binding domain [12] and upstream of the putative Rab8a binding domain. This mutation was predicted to truncate the protein by almost half (Fig. 1A) and is near a similar mutation in humans (p.Gln1265X) that associated with LCA and JBTS (https://cep290base.cmgg.be/). We confirmed the mutation by direct sequencing (Fig. 1B). To date, no paralogue to *cep290* has been reported in zebrafish. To assess the impact of the *fh297* allele on gene expression, *cep290* mRNA was quantified by digital droplet PCR (ddPCR). In adult animals, retinal expression of *cep290* mRNA was reduced by 55% in mutants compared to expression in wild-type siblings (Fig. 1C). Efforts to measure Cep290 protein by western blot were unsuccessful, despite multiple attempts with both commercial [41] and custom designed antibodies [42]. In fibroblasts derived from an LCA patient with the c.2991+1665A>G mutation, wild-type *CEP290* transcripts were similarly reduced by ∼60%, which resulted in a corresponding reduction in protein levels by ∼80% [43].

**Figure 1.**
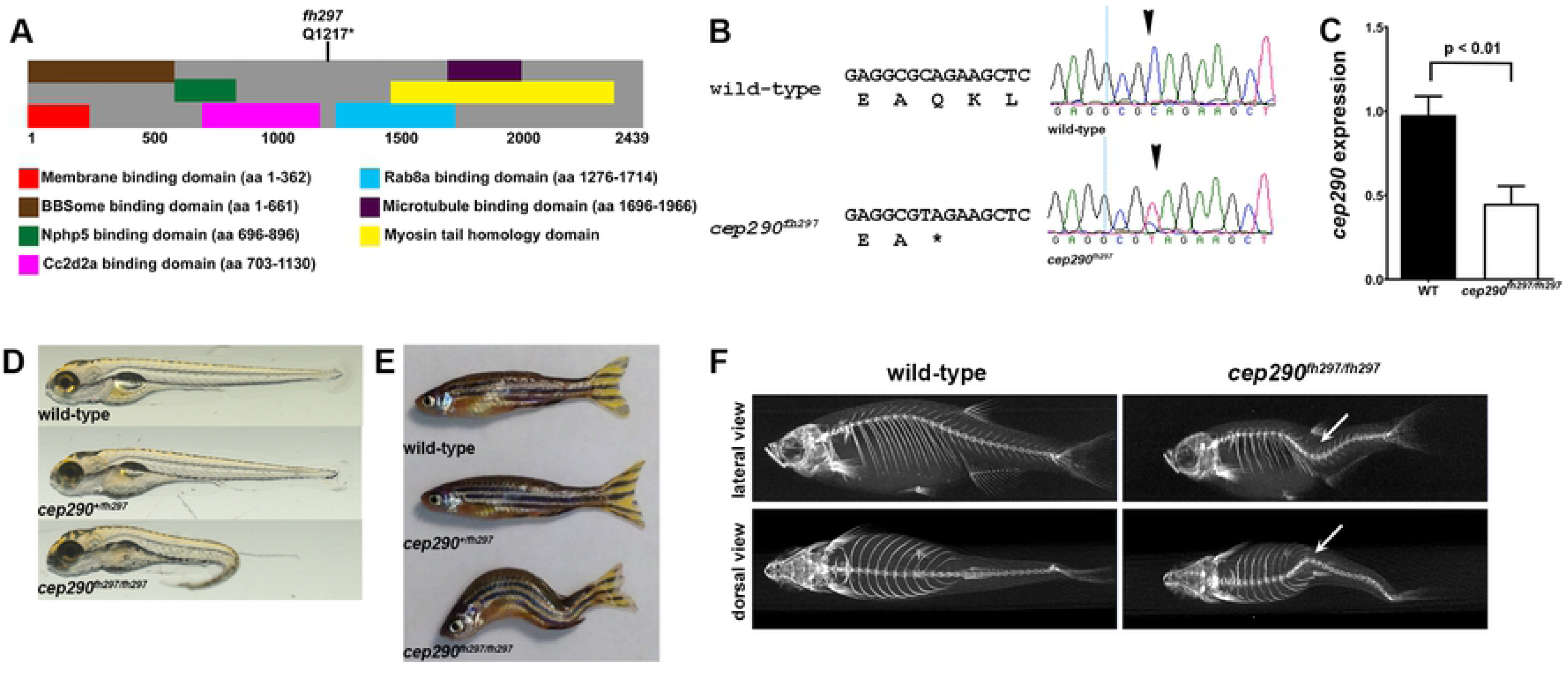
Genetic mapping and identification of *cep290* mutant allele. (A) Schematic structure of zebrafish Cep290 illustrating the location of predicted protein domains. Domain structure is based on prior results [12, 44, 45]. The *cep290*^*fh297*^ allele generates a premature stop codon at amino acid 1217. (B) Chromatograms of Sanger sequencing reactions of cDNAs from wild-type and homozygous *cep290*^*fh297/fh297*^ mutant confirming the C>T replacement. (C) Quantification of *cep290* mRNA in 6 month old wild-type and mutant retinas by digital droplet PCR (ddPCR). (D) Lateral views of representative wild-type (top), heterozygous (middle), and homozygous (bottom) mutants at 5 dpf. At larval stage 30% of *cep290*^*fh297/fh297*^ mutant animals show ventral tail curvature. (E) Lateral view of 7 month old wild-type, heterozygous and a representative *cep290*^*fh297/fh297*^ mutants displaying distorted vertebral column. At adult stage 100% of the homozygous mutant animals show scoliosis of the vertebral column. (F) Representative μCT-generated images of lateral (top) and dorsal (bottom) views of adult wild-type and *cep290*^*fh297/fh297*^ mutants. Images demonstrate that spinal curvature deviates within the dorsal/ventral plane as well as curves laterally (arrows).

At 5 days post fertilization (dpf) *cep290*^*fh297/fh297*^ mutants exhibited a sigmoidal tail curvature and did not yet have an inflated swim bladder (Fig. 1D). These phenotypes were not fully penetrant with only 29% of mutant larvae exhibiting such characteristics (17 of 58 confirmed *cep290*^*fh297/fh297*^ mutants). Furthermore, these phenotypes were similar to, but distinct from phenotypes of other zebrafish mutants with defective cilia formation, such as *ift88* or *cc2d2a* mutants, which exhibit a ventral axis curvature [40, 46, 47]. All *cep290*^*fh297/fh297*^ mutants exhibited normal otolith numbers and only 6.8% (4 of 58 confirmed *cep290*^*fh297/fh297*^ mutants) developed kidney cysts by 5 dpf. Although *cep290*^*fh297/fh297*^ mutants routinely survived to adulthood in Mendelian ratios, approximately 17% fewer *cep290*^*fh297/fh297*^ mutants were present at 12 months of age than would be expected (25 out of 120 total fish). The *cep290*^*fh297/fh297*^ mutants were unable to breed and 100% of the mutants exhibit a severe scoliosis (Fig. 1E, F), a phenotype previously linked to defective cilia [48] and a pathology has been reported in a a subset of Joubert Syndrome patients with *CEP290* mutations [49]. The abnormal spinal curvature was also assessed by micro-computed tomography (μCT) and revealed a significant deviation in spinal curvature that was pronounced in both the dorsal-ventral axis as well as a lateral curvature (Fig. 1F).

As the *cep290*^*fh297*^ allele encodes a nonsense mutation, we were curious about the relatively mild phenotype compared to the *Cep290* mouse knockout models [14]. A recent hypothesis proposed that if nonsense mutations occur in exons that begin and end in the same reading frame, those exons can be preferentially skipped in a process known as “nonsense-induced alternative splicing” [50]. These alternatively spliced transcripts can escape nonsense-mediated decay and produce near-full-length protein. Such phenomenon were reported to occur in Leber Congenital Amaurosis and Senor-Løken Syndrome patients with mutations in *CEP290* [25, 37]. The *cep290*^*fh297*^ allele is a nonsense mutation occurring in exon 29 and exon skipping would maintain the reading frame. We performed RT-PCR on cDNA from adult retinas from wild-type and *cep290* mutant retinas and used primers that spanned the mutated exon. We showed that the mutant exon was present in all detectable transcripts, indicating that nonsense-mediated alternative splicing did not occur for this mutation in zebrafish (Figs. 2A, B).

**Figure 2.**
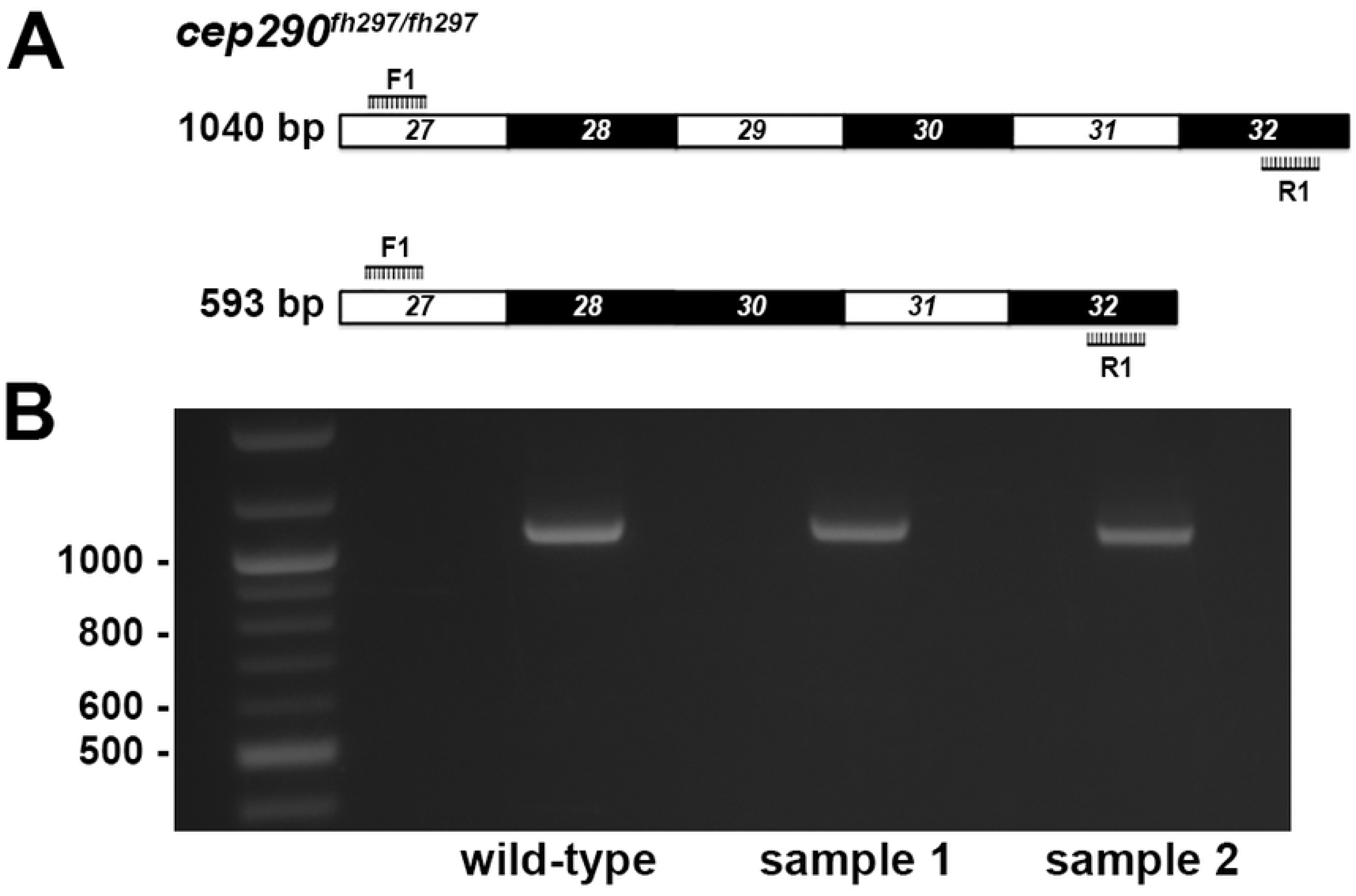
Exon skipping of *cep290* exons 29 does not occur in zebrafish retinas. (A) Forward (F1) and reverse (R1) PCR primers were designed to bind sequences in exons 27 and 32 in order to flank exon 29 harboring the mutant *cep290*^*fh297*^ allele. Exon skipping would result in a truncated PCR product of 593 bp, while retention of exon 29 would result in a full-length 1040 bp product. (B) Results of PCR reactions from cDNAs generated from wild-type and samples of two individual mutants resulted in full-length products of 1040 bp. 100 bp ladder shown in lane 1.

### Functional vision is preserved in *cep290* mutant larvae

As humans with *CEP290* mutations report variable loss of visual function [9], we asked whether the visual performance of zebrafish *cep290* mutants was also compromised. Visual acuity is a measure of the spatial resolving power of the visual system and is mainly driven by cones [51, 52]. Larval zebrafish visual function can be readily assessed using the optokinetic response (OKR) assay by presenting larvae with a moving grating stimuli that varies in either contrast or spatial frequency [39]. Detecting contrast differences of a stimulus presented at fixed spatial and temporal frequencies is a general method of testing function vision, while detecting the changes in spatial frequency of a stimulus at a fixed contrast under bright illumination assesses cone acuity. Because larval zebrafish rely exclusively on cone function at 5-6 dpf [53, 54], all recordings were done under photopic conditions [55]. We measured the OKR gain, which is defined by the ratio between stimulus velocity and eye velocity [38-40], of wild-type, and *cep290*^*fh297/fh297*^ mutants between 5-6 dpf using established parameters [39, 56] and reproduced by our laboratory [38, 40]. Wild-type larvae (n = 12) showed a linear relationship between gain and the logarithm of contrast (Fig. 3A). Interestingly, the *cep290*^*fh297/fh297*^ mutants (n = 26) exhibited normal OKR responses to changes in stimulus contrast and spatial frequency (Fig. 3A, B).

**Figure 3.**
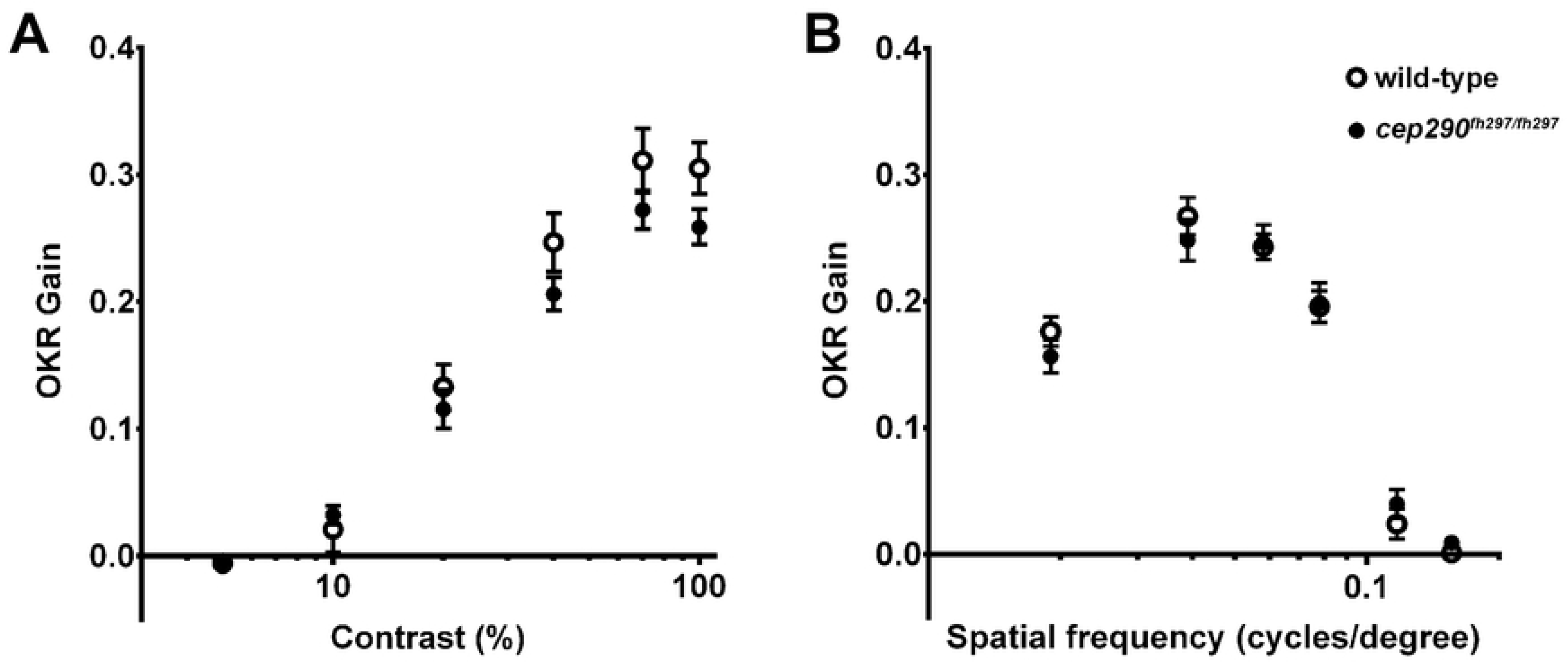
Visual performance is not affected in *cep290* mutant larvae at 5 dpf. (A) Optokinetic response (OKR) contrast response function of 5 dpf wild-type larvae (n = 11; open circles) and *cep290*^*fh297/fh297*^ mutants (n = 26; closed circles). No significant differences were found. (B) OKR spatial frequency results for wild-type larvae (n = 13) and *cep290*^*fh297/fh297*^ mutants (n = 15). Error bars = s.e.m.

### Photoreceptor degeneration in *cep290*^*fh297/fh297*^ mutants

We next examined the retinal anatomy of wild-type and *cep290*^*fh297/fh297*^ mutants by light microscopy at 5 dpf, 3 months post fertilization (mpf), 6 mpf, and 12 mpf (Fig. 4). Normal retinal lamination and cellular differentiation was observed in *cep290*^*fh297/fh297*^ mutants at 5 dpf (Fig. 4A), indicating that retinal development did not require Cep290. At 3 mpf, we noticed fewer and more disorganized cone outer segments in *cep290*^*fh297/fh297*^ mutants (Figs. 4B, white arrow). Cone disorganization in *cep290*^*fh297/fh297*^ mutants was progressive and by 6 mpf the loss of cone outer segments (COS) was more evident (Figs. 4C). By 12 mpf, only a few discernable cones remained in the *cep290*^*fh297/fh297*^ mutants (Figs. 4D, arrows). Also noticeable was the continued thinning of the retinal outer nuclear layer (ONL) in *cep290*^*fh297/fh297*^ mutant retinas when compared to wild-type animals (Figs. 4E, F). The thickness of the ONL was reduced across the peripheral and central retina, although the difference was only statistically significant in the dorsal retina (Fig. 4E). When the rows of nuclei in the ONL was quantified, a statistically significant difference was seen in the periphery of the dorsal retina (Fig. 4F; 4.1±0.3 vs 3.1±0.3 rows of nuclei, *P* < 0.05; n = 6).

**Figure 4.**
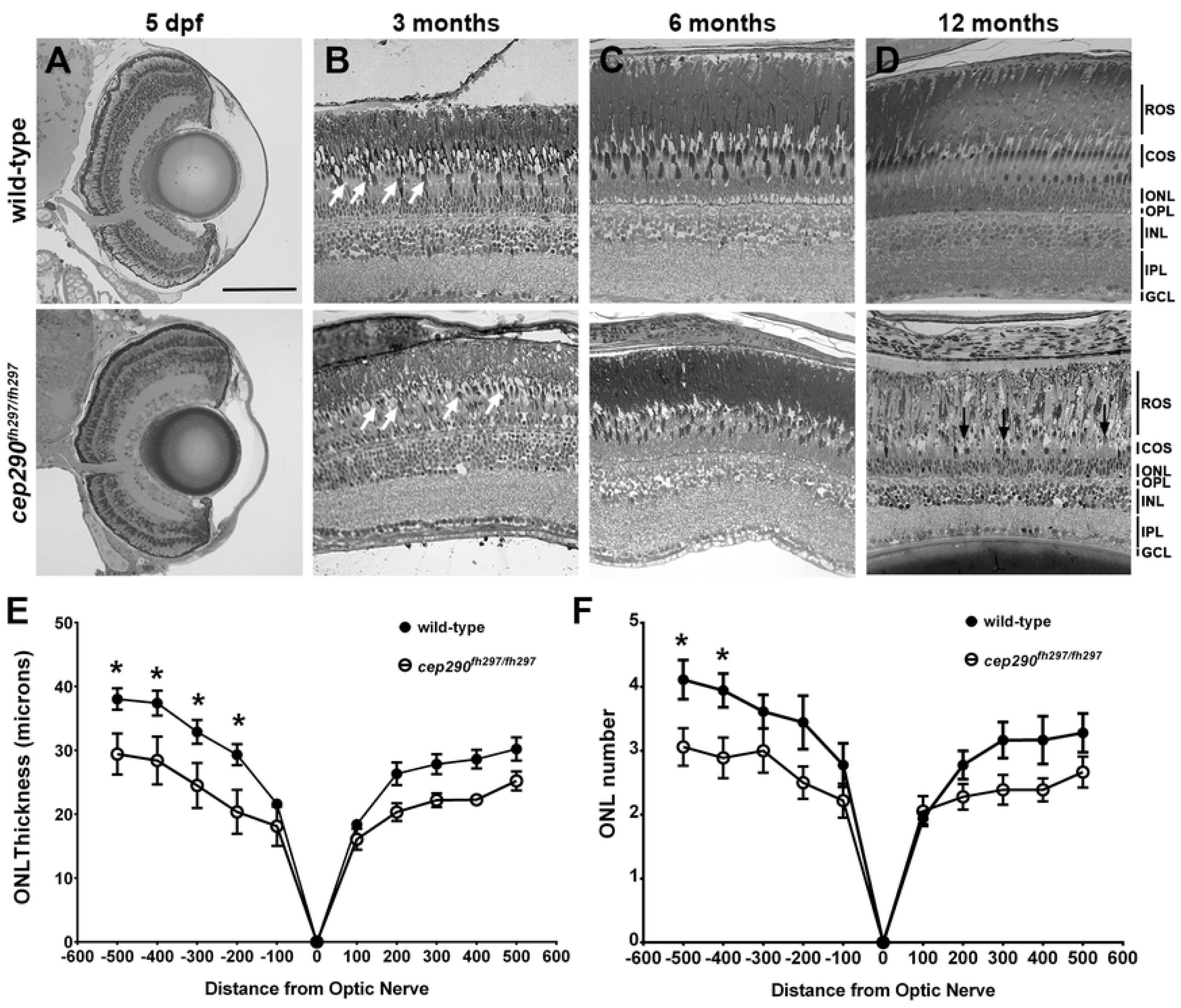
Progressive cone loss in adult *cep290*^*fh297/fh297*^ mutants. Methylene blue stained transverse histological sections of retinas from wild-type (top) and *cep290*^*fh297/fh297*^ mutants (bottom) at 5 dpf (A); 3 months of age (B), 6 months of age (C), and 12 months of age (D). At 3 months, the *cep290*^*fh297/fh297*^ mutants (bottom) had noticeably fewer cones (white arrows) and thinning of the cone outer segment (COS) layer. Few cones were observed at 12 months of age in *cep290*^*fh297/fh297*^ mutants (black arrows). (E) Quantification of ONL thickness and (F) rows of nuclei in the ONL at different distances from the optic nerve in both the dorsal (negative numbers; left) and ventral (positive numbers; right) retina of *cep290*^*fh297/fh297*^ mutants and wild-type sibling controls at 8 months of age. Data are shown as means ± SEM (n = 6, \**P* ≤ 0.05). ROS, rod outer segments; COS, cone outer segments; ONL, outer nuclear layer; OPL, outer plexiform layer; INL, inner nuclear layer; IPL, inner plexiform layer; GCL, ganglion cell layer. Scale bar: 100 μm

We next used electron microscopy to examine retinal sections of wild-type and *cep290*^*fh297/fh297*^ mutant adults (6 mpf) to determine how loss of Cep290 affected photoreceptor structure. In *cep290*^*fh297/fh297*^ mutants, few cone outer segments were observed and the outer retina of *cep290*^*fh297/fh297*^ mutants was more disorganized than that seen in wild-type retinas (Figs. 5A-C). At higher magnification, whereas the outer segments of wild-type animals contained highly organized stacks of disc membranes, the disc membranes were fragmented and the outer segments appear to be disintegrating in *cep290*^*fh297/fh297*^ mutants (Figs. 5D, E). We did not, however, observe any consistent accumulation of vesicular material adjacent to the ciliary base or other signs of disrupted ciliary trafficking (Figs. 5D, E; white arrowheads). These results suggest that loss of Cep290 disrupts cone outer segment structure and causes cell death.

**Figure 5.**
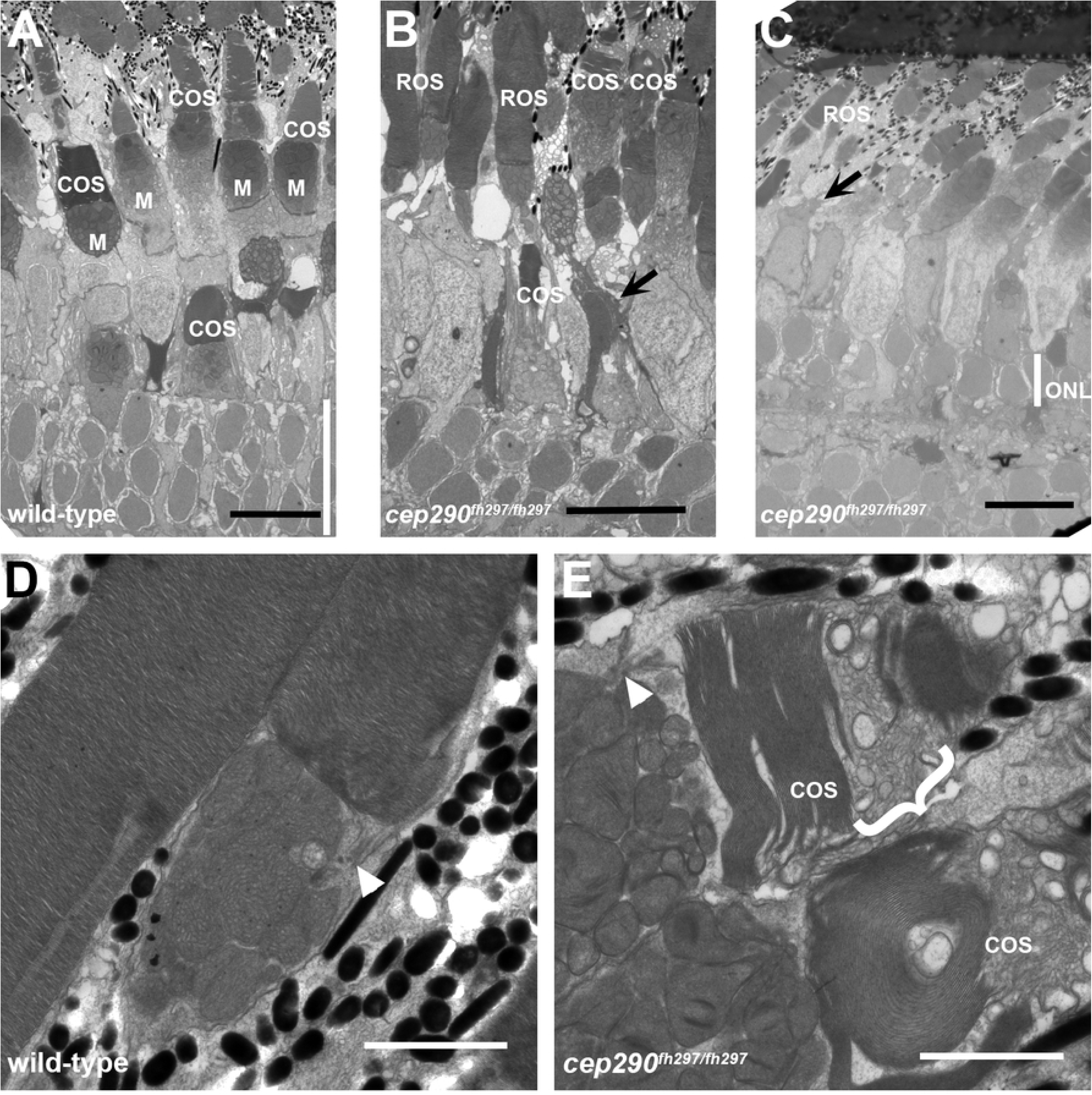
Cone degeneration marked by outer segment disorganization in *cep290*^*fh297/fh29*^ mutants. (A-C) Transmission electron micrographs of retinal sections from 6 month old wild-type (A) and *cep290*^*fh297/fh297*^ mutant adults (B, C). Cone outer segments and mitochondria in the ellipsoids are visible in the wild-type retina. In the *cep290*^*fh297/fh297*^ mutant retinas, cone outer segments are missing or disorganized (arrows) and the outer nuclear layer (ONL, white line) is reduced to 1-2 nuclei. (D, E) At higher magnification, the outer segment disc membranes are tightly stacked in wild-type retinas. In *cep290*^*fh297/fh297*^ mutants, numerous vesicular structures and disorganized membranes are seen in cone outer segments (bracket). Rod outer segments are largely preserved and the connecting cilia are shown (white arrowheads). Scale bars: 10 µm (A-C); 2 µm (D, E).

### Progressive cone degeneration in *cep290*^*fh297/fh297*^ mutants

To track the progression of photoreceptor degeneration, immunohistochemistry was performed on retinal cryosections of *cep290*^*fh297/fh297*^ mutants at 3, 6, and 12 months of age. Retinas were stained with peanut agglutinin lectin (PNA) to label the interphotoreceptor matrix surrounding cone outer segments [57] and Zpr-1, a monoclonal antibody that recognizes arrestin-3 like (Arr3L) on the cell bodies of red- and green-sensitive double cones [58, 59]. Similar to the results from plastic histology, the PNA-labeled cone outer segments were less organized and the inner segments appeared less densely packed in the *cep290*^*fh297/fh297*^ mutants at 3 months of age as compared to wild-type siblings (Figs. 6A-F). Cone degeneration in the *cep290*^*fh297/fh297*^ mutants was more apparent by 6 months of age (Figs. 6G-L) and by 12 months of age, very few cones remained (Figs. 6M-R). In older *cep290*^*fh297/fh297*^ mutants, some cones had Zpr-1 positive inner segments but lacked PNA-positive outer segments (Figs. 6L, R; arrows), indicating that outer segment loss preceded cone death. This is consistent with a role for Cep290 in sensory cilia maintenance. The density of PNA-positive cone outer segments in wild-type and *cep290*^*fh297/fh297*^ mutants were quantified at each time point (Fig. 7). The results indicated that cone degeneration initiated by 3 months of age in *cep290*^*fh297/fh297*^ mutants and progressed such that very few cone outer segments (1.6 COS / 50 μm) remained by 12 months of age.

**Figure 6.**
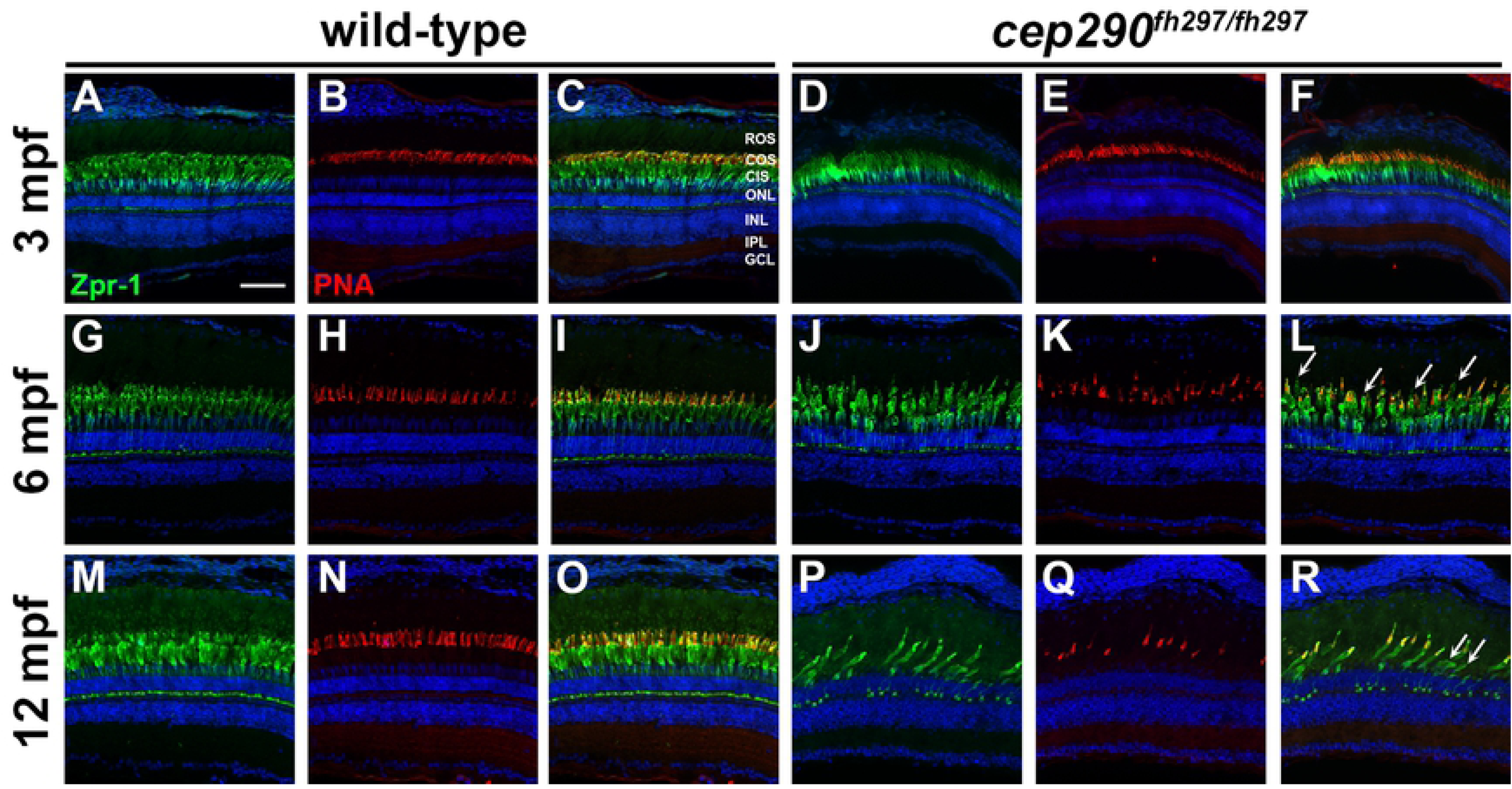
Cone outer segment degeneration progresses with age in *cep290*^*fh297/fh297*^ mutants. Immunohistochemistry of retinal cryosections stained with peanut agglutinin (PNA) to label cone outer segments and Zpr-1 (green) to label red/green double cones of wild-type and *cep290*^*fh297/fh297*^ mutants. Views from dorsal retinas are shown. (A-F) Retinas from 3-month old adults. (G-L) Retinas from 6-month old adults. (M-R) Retinas from 12-month old adults. Arrows denote cones that were negative for PNA but positive for Zpr-1. ROS, rod outer segments; COS, cone outer segments; CIS, cone inner segments; ONL, outer nuclear layer; INL, inner nuclear layer; IPL, inner plexiform layer; GCL, ganglion cell layer. Scale bar: 50 μm

**Figure 7.**
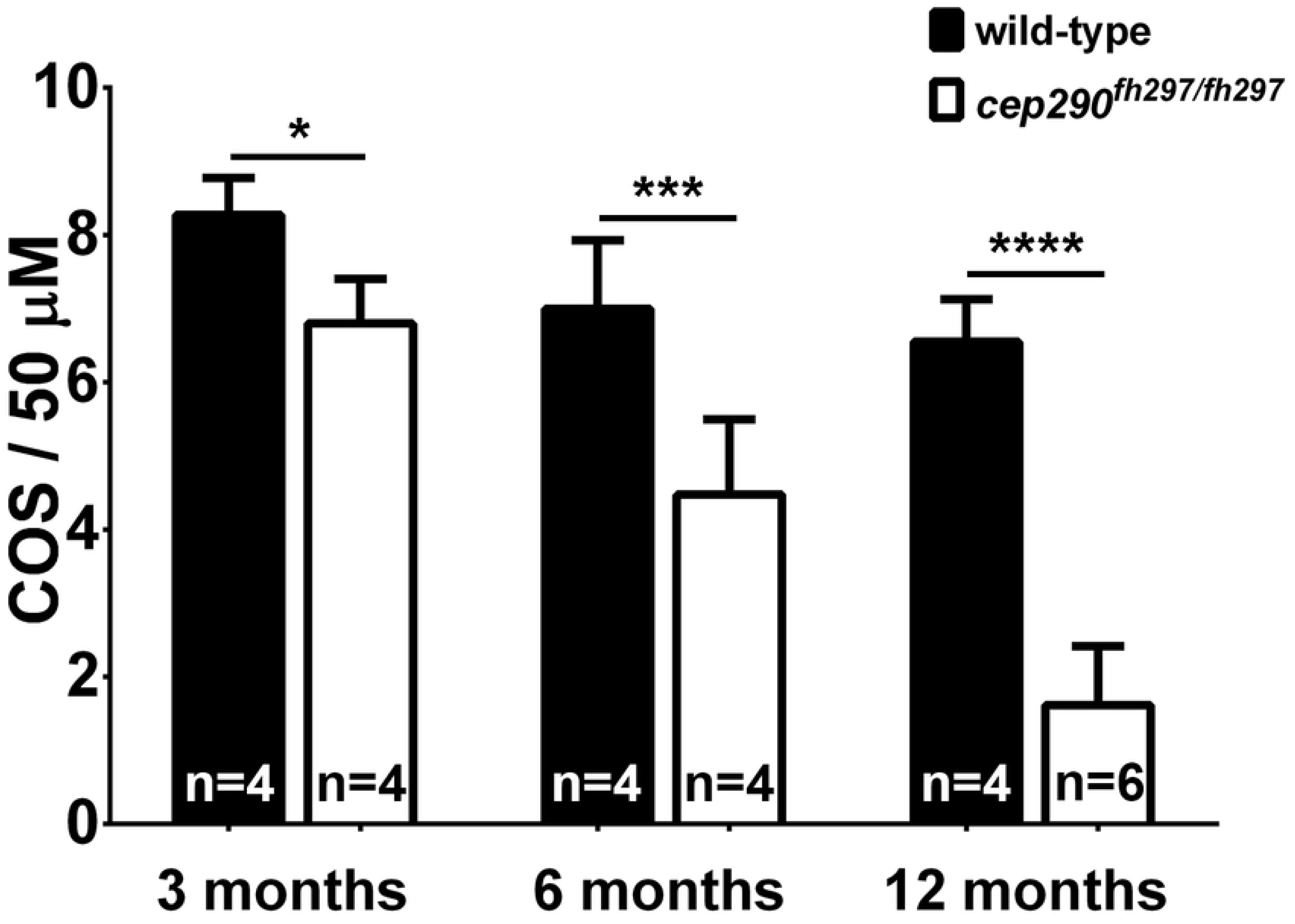
Cone outer segment density declines with age in *cep290*^*fh297/fh297*^ mutants. Quantification of cone outer segment density at ages from Figure 6. The number of independent fish used for each measurement is indicated. \**P* < 0.05; *** *P* < 0.0005; **** *P* < 0.0001 as determined by a 2-way ANOVA with a Multiple Comparisons test and Sidek corrections.

### Distribution of rod outer segment proteins in *cep290*^*fh297/fh297*^ mutants

Loss of Cep290 is associated with rapid loss of rods and rhodopsin mislocalization in the mouse *cep290*^*rd16/rd16*^ mutant [20]. Rhodopsin is a G-protein coupled receptor with seven transmembrane domains that passes along the ciliary plasma membrane before becoming incorporated into disc membranes within the outer segment. To evaluate the effects of Cep290 loss on rods in zebrafish, we stained retinal cryosections from *cep290*^*fh297/fh297*^ mutants and wild-type siblings with the monoclonal antibody Zpr3, which recognizes rhodopsin. At 3 mpf, when the first signs of cone degeneration were observed in *cep290*^*fh297/fh297*^ mutants, rhodopsin localized to the outer segments of both *cep290*^*fh297/fh297*^ mutants and wild-type control animals (Figs. 8A-D). By 6 mpf, when cone degeneration was pronounced, a significant amount of rhodopsin mislocalized to the inner segments and cell bodies (Figs. 8E-H; arrowheads). Rhodopsin continued to be mislocalized at 12mpf (Figs. 8I-L; arrowheads), but significant loss of rod outer segment material was not observed. At 12 mpf, however, we noticed that the gap between the rod outer segments and the ONL, which typically is occupied by cone nuclei, was considerably smaller in 12 month-old animals as compared to wild-type siblings (Figs. 8I, J; white brackets).

**Figure 8.**
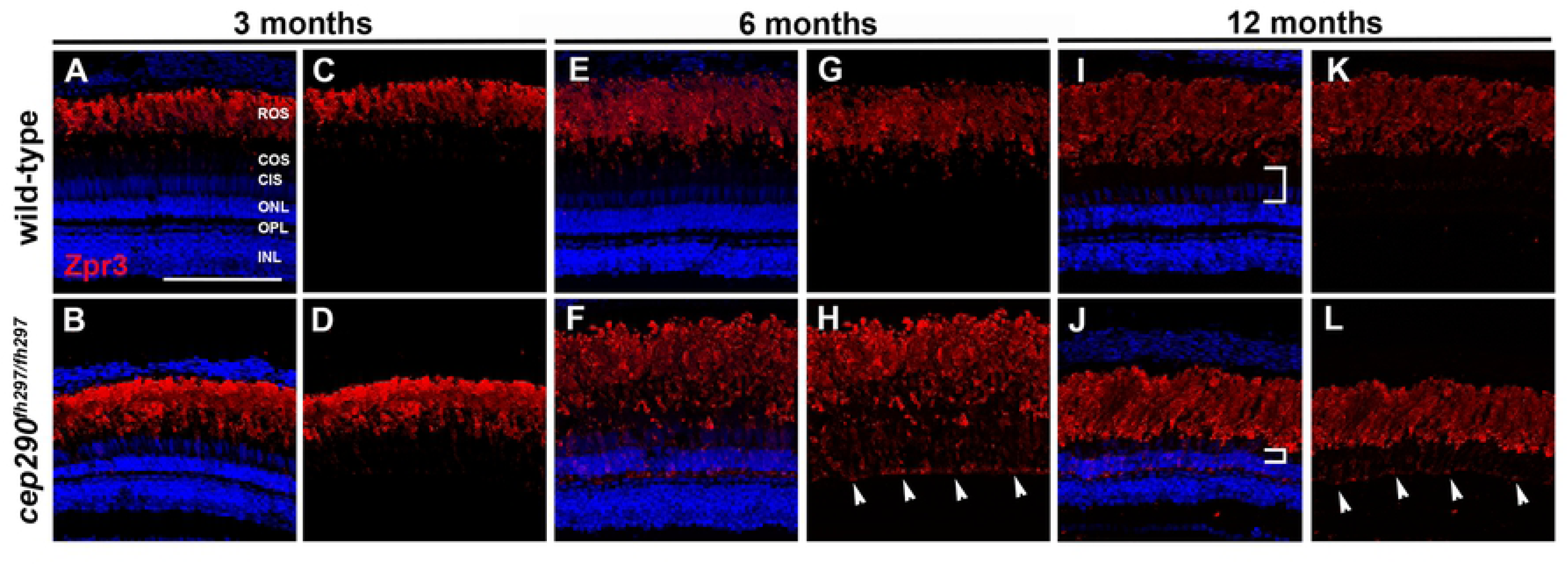
Mislocalization of rhodopsin in *cep290*^*fh297/fh297*^ mutants. (A-D) Images show cryosections labeled with Zpr3 (red) to mark rhodopsin and DAPI (blue) to label nuclei in the dorsal retinas from *cep290*^*fh297/fh297*^ mutants and wild-type siblings at 3 months of age; (E-H) 6 months of age, and (I-L) 12 months of age. At later ages, the distance between the base of the rod outer segments and the outer nuclear layer decreases due to loss of cone nuclei (I, J; white brackets). Arrowheads note rhodopsin mislocalization. ROS, rod outer segments; COS, cone outer segments; CIS, cone inner segments; ONL, outer nuclear layer; OPL, outer plexiform layer; INL, inner nuclear layer. Scale bar: 100 μm

Active transport of cytosolic and transmembrane proteins (e.g. rhodopsin) through the ciliary TZ to the photoreceptor outer segments requires intraflagellar transport (IFT), while transport of lipidated protein cargo requires a distinct targeting system utilizing either PDE6D or UNC119 [60]. We therefore investigated the localization rhodopsin kinase (GRK1) and the β subunit of rod transducin (GNB1), to determine if loss of Cep290 also disrupts trafficking of lipidated ciliary proteins. GRK1 is a prenylated membrane protein that requires the function of Retinitis Pigmentosa 2 (RP2) for proper transport to rod outer segments [61]. Membrane association of GNB1 requires direct binding to the protein RP2 and loss of RP2 disrupts outer segment trafficking of both GNB1, GRK1, and other prenylated proteins [62-64]. In retinal cryosections from animals at both 6 months and 12 months of age, we found that GRK1 localized to the rod outer segments of *cep290*^*fh297/fh297*^ mutants, similar to wild-type siblings (Figs. 9A-H). At both 6 and 12 months of age, the majority of GNB1 also localized to the rod outer segments of both wild-type and *cep290*^*fh297/fh297*^ mutants (Figs. 9I-P). Taken together, these results suggest that loss of Cep290 specifically affects rhodopsin localization without broadly impairing transport of all outer segment proteins.

**Figure 9.**
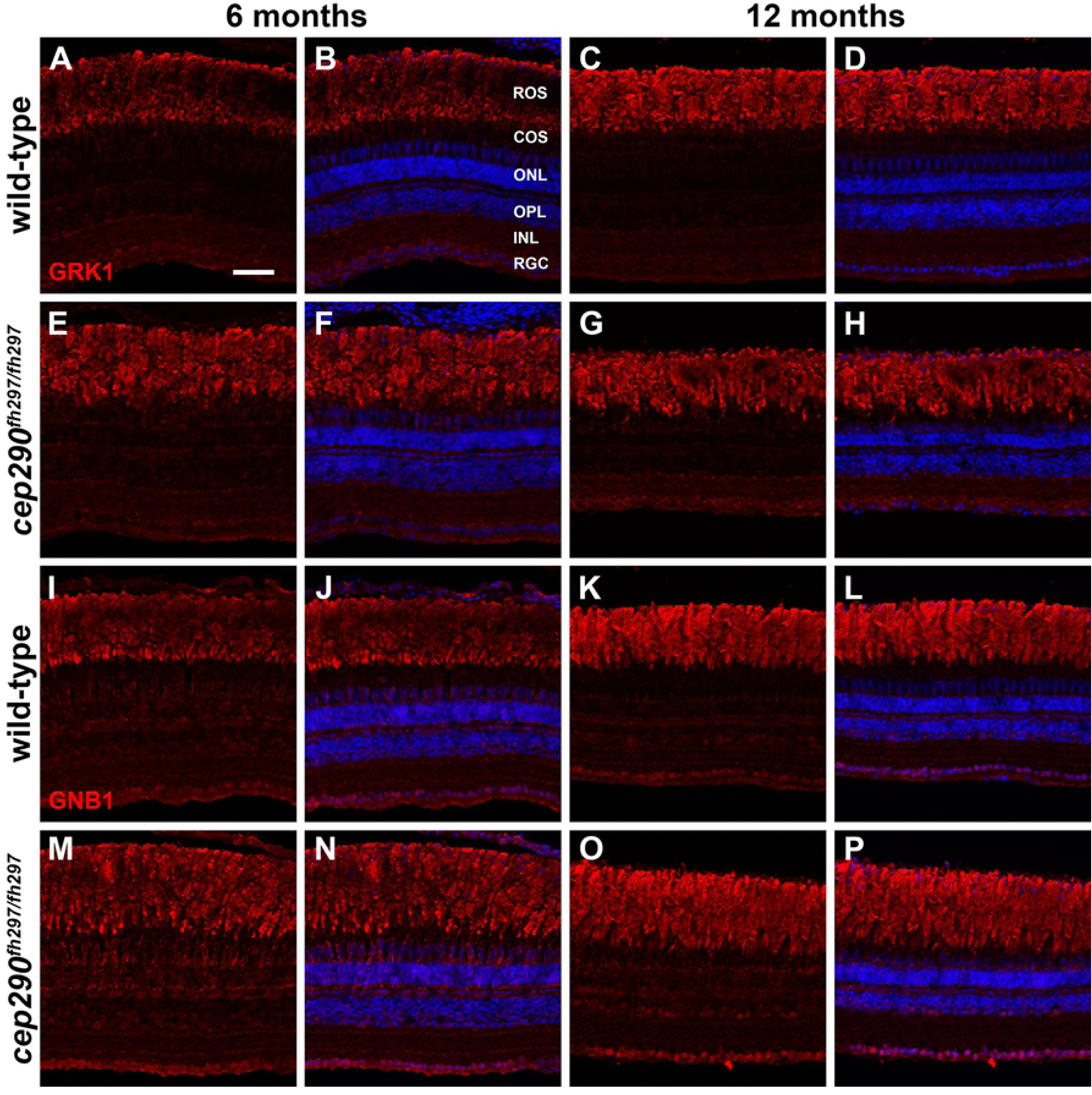
Immunolocalization of GRK1 and GNB1 in *cep290*^*fh297/fh297*^ mutants at 6 and 12 months of age. Retinal cryosections of *cep290*^*fh297/fh297*^ mutants and wild-type siblings were stained with polyclonal antibodies against rhodopsin kinase (A-H; GRK1) or with GNB1 polyclonal antibodies against the β subunit of rod transducin (I-P) to label rod outer segments at both 6 and 12 months of age. ROS, rod outer segments; COS, cone outer segments; ONL, outer nuclear layer; OPL, outer plexiform layer; INL, inner nuclear layer; RGC, retinal ganglion cells. Scale bar: 50 μm

In response to retinal injury, zebrafish typically exhibit a robust capability of regenerating lost neurons, including photoreceptors [65, 66]. In the uninjured retina, Müller glia in the inner nuclear layer (INL) will periodically divide and produce unipotent rod progenitor cells that migrate to the ONL where they can proliferate as rod precursors and differentiate into rod photoreceptors [67-69]. In response to acute retinal damage, however, the Müller glia will dedifferentiate, undergo cellular reprogramming, and produce multipotent retinal progenitors that proliferate and are able to differentiate into all retinal cell types, including cones [66, 70]. Given this regenerative capacity, it was surprising to observe photoreceptor degeneration *cep290*^*fh297/fh297*^ mutants. To determine if *cep290*^*fh297/fh297*^ mutants attempted regeneration, retinas from 3-month and 6-month old *cep290*^*fh297/fh297*^ mutants and wild-type siblings were stained with antibodies against proliferating cell nuclear antigen (PCNA), which is a marker of cell proliferation, and quantified the number of PCNA+ cells in the ONL and inner nuclear layer (INL). Only a small number of individual proliferating cells were seen in the INL of 3 month old *cep290*^*fh297/fh297*^ mutants (7.7±3.5) or wild-type siblings (10.9±1.5) and no statistical difference was found (Figs. 10A, B; n = 6; *P* = 0.15). Compared to the INL, there were up to 10-fold more proliferating cells in the ONL of 3-month old mutant (82.7±17.2) and wild-type siblings (62.3±12.8), but still no statistical difference seen (Figs. 10A, C; n = 6; *P* = 0.06). At 6 months of age, however, significantly more proliferating cells were found in both the INL and ONL of *cep290*^*fh297/fh297*^ mutants (Figs. 10B, C). Compared to wild-type siblings, there were 3-fold more proliferating cells in the INL and 10-fold more cells in the ONL of *cep290*^*fh297/fh297*^ mutants. Of note, proliferating cells were 10-fold more abundant in the ONL than in the INL of *cep290*^*fh297/fh297*^ mutants (note differences in Y-axes in Figs. 10B, C). This suggests photoreceptor degeneration in *cep290*^*fh297/fh297*^ mutants triggers robust proliferation of rod progenitor cells but only limited proliferation of Müller glia.

**Figure 10.**
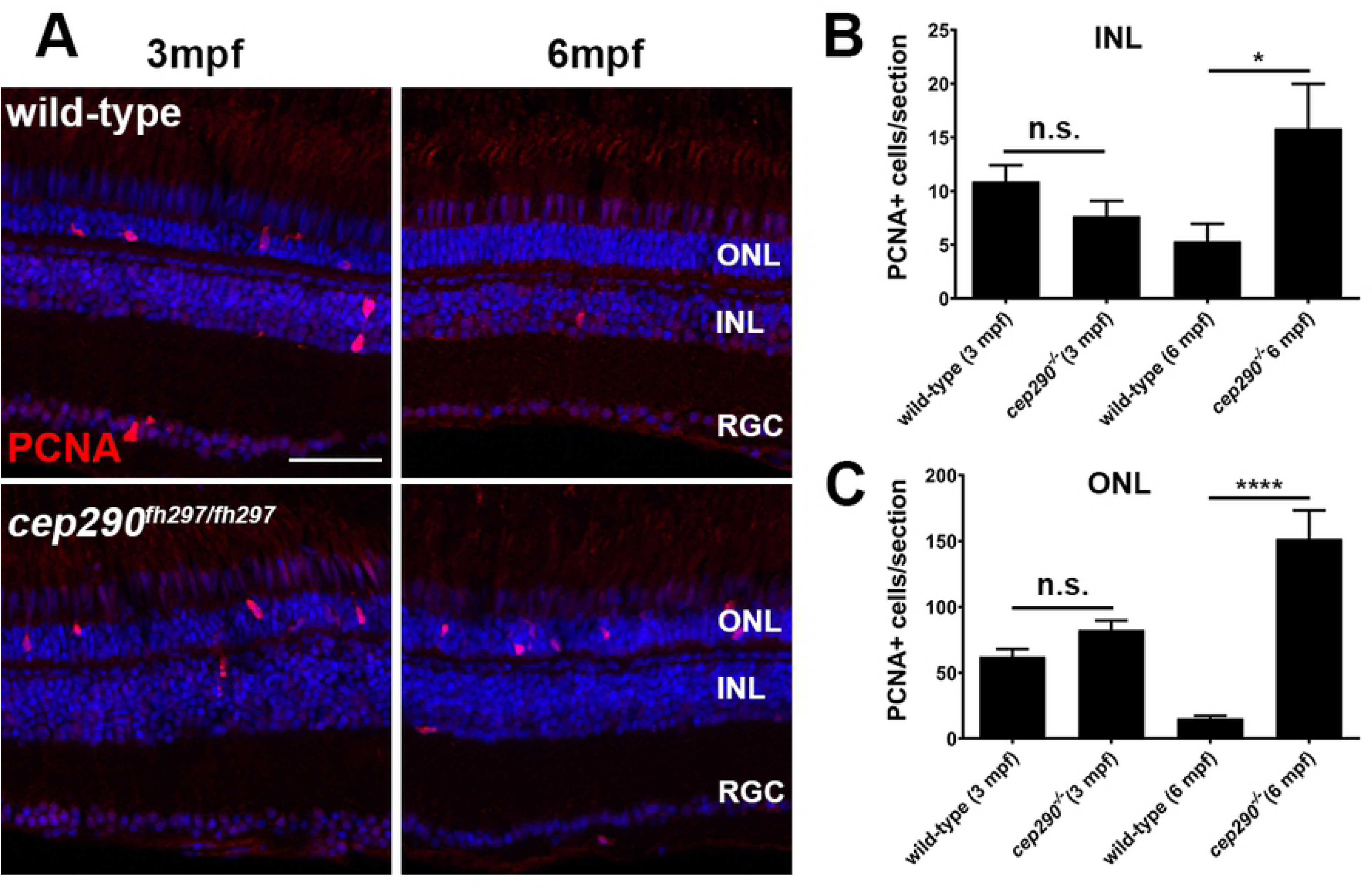
PCNA localization in *cep290*^*fh297/fh297*^ mutants at 3 and 6 months of age. (A) PCNA immunolocalization in cryosections of the dorsal retina of wild-type (top) and *cep290*^*fh297/fh297*^ mutants (bottom) at 3-months and 6-months of age. (B) PCNA positive cells were quantified in the INL from cryosections of the both dorsal and ventral retina at different ages. (C) Quantification of PCNA in the ONL from cryosections across the dorsal and ventral retina at different ages. A significant increase in PCNA immunoreactivity was seen in both the INL and ONL of *cep290*^*fh297/fh297*^ mutants at 6 months of age. Quantification was performed on cryosections of individual retinas from *cep290*^*fh297/fh297*^ mutants (n = 6) and wild-type siblings (n = 5) at the stated ages. Values represent the mean ± s.e.m. \**P* < 0.05; **** *P* < 0.0001 as determined by an unpaired t-test. ONL, outer nuclear layer; INL, inner nuclear layer; RGC, retinal ganglion cells. Scale bar: 50 μm

### Effects of the combined loss of *cep290* and cilia genes *ahi1*, *cc2d2a*, or *arl13b* differentially affects visual acuity but does not exacerbate photoreceptor degeneration

A leading hypothesis to explain phenotypic variability in ciliopathies is the effects of mutations in second-site modifiers [26-29]. Typically, heterozygosity (i.e. loss of one allele) in cilia genes exacerbates the phenotypes observed in homozygous mutants of other cilia genes. Analysis of animals with homozygous mutations in two distinct genes may reflect the additive effect of two independent phenotypes and not necessarily a role for second-site modifiers. Because the loss of *cep290* leads to slow cone degeneration in zebrafish, we asked whether heterozygous mutations in genes encoding other cilia proteins would accelerate degeneration. The Cc2d2a and Ahi1 proteins are components of the MKS module that make up part of the TZ [28]. In humans, mutations in *AHI1* and *CC2D2A* cause Joubert Syndrome and both genes have been proposed as potential modifiers of *CEP290* [12, 33]. Cep290 directly binds Cc2d2a through an N-terminal domain of Cep290 [12] and Cep290 is predicted to bind Ahi1, suggesting that these connections are critical for TZ assembly or stability. Mutations in *ARL13B* also result in Joubert Syndrome. The Arl13b protein is required for axoneme extension and photoreceptor outer segment formation [34]. Importantly, the zebrafish *cc2d2a*^*-/-*^ and *ahi1*^*-/-*^ mutants show defects in photoreceptor OS structure during larval stages [40, 47] while the *arl13b*^*-/-*^ zebrafish mutant undergoes a progressive photoreceptor degeneration [35]. The known and proposed biochemical and genetic interactions, as well as similar protein localization patterns in the transition zone and axoneme, suggested that these genes could potentially modulate phenotypes associated with Cep290 mutations.

As *cep290*^*fh297/fh297*^ adults were unable to breed naturally, heterozygous animals were mated to generate *cep290*^*+/fh297*^;*ahi1*^*+/-*^, *cep290*^*+/fh297*^;*cc2d2a*^*+/-*^, and *cep290*^*+/fh297*^;*arl13b*^*+/-*^ adults. Pairwise crosses from these adults generated all possible genotypes in the expected Mendelian ratios. The double homozygous mutants (e.g. *cep290*^*fh297/fh297*^;*ahi1*^*-/-*^ and *cep290*^*fh297/fh297*^;*cc2d2a*^*-/-*^; and *cep290*^*fh297/fh297*^;*arl13b*^*-/-*^) did not survive beyond 14 dpf. The *cep290*^*fh297/fh297*^;*ahi1*^*+/-*^, *cep290*^*fh297/fh297*^;*cc2d2a*^*+/-*^; and *cep290*^*fh297/fh297*^;*arl13b*^*+/-*^ mutants were viable beyond 12 months and were indistinguishable from *cep290*^*fh297/fh297*^ mutants.

We next wanted to determine if loss of *cep290* sensitizes photoreceptors to the additional loss of one allele of either *ahi1, arl13b,* or *cc2d2a* and would accelerate cone or rod degeneration. We first assessed cone degeneration by immunohistochemistry using the markers PNA and Zpr-1 on retinas from 6-month old *cep290*^*fh297/fh297*^;*ahi1*^*+/-*^ (Fig. 11A), *cep290*^*fh297/fh297*^;*cc2d2a*^*+/-*^ mutants (Fig. 12A) and *cep290*^*fh297/fh297*^;*arl13b*^*+/-*^ mutants (Fig. 13A) when compared to wild-type and *cep290*^*fh297/fh297*^ mutants. We quantified cone density within the dorsal retina for each genotype (Figs. 11D, 12D, 13D). Whereas *cep290*^*fh297/fh297*^ mutants exhibited reduced numbers of cone outer segments, no additional increase in cone loss was observed in the *cep290*^*fh297/fh297*^;*ahi1*^*+/-*^, *cep290*^*fh297/fh297*^;*cc2d2a*^*+/-*^ or *cep290*^*fh297/fh297*^;*arl13b*^*+/-*^ mutants (Figs. 11D, 12D, 13D). Retinal sections were also stained with antibodies against rhodopsin and rhodopsin kinase (GRK1) to assess trafficking of rod outer segment proteins (Figs. 11B, 11C, 12B, 12C, 13B, and 13C). Rhodopsin mislocalization was quantified by measuring integrated fluorescence density in the rod inner and outer segments (see Methods). Whereas considerable rhodopsin mislocalization was observed in *cep290*^*fh297/fh297*^ mutants, there was no significant exacerbation of this phenotype by the additional loss of one allele of *ahi1, cc2d2a*, or *arl13b* (Figs. 11E, 12E, 13E). GRK1 localized to the rod outer segments in wild-type animals and in all mutant genotypes (Figs. 11C, 12C, 13C).

**Figure 11.**
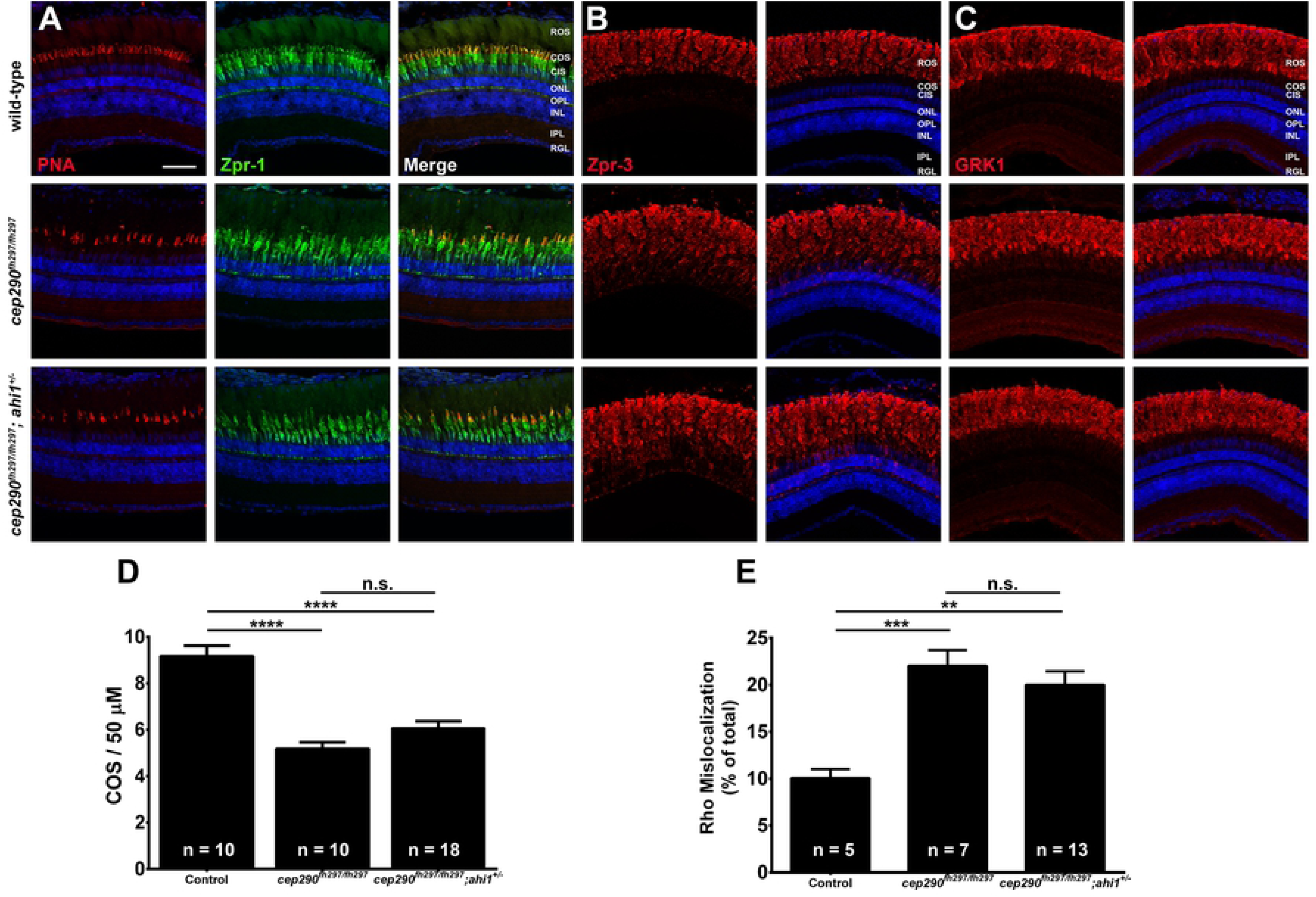
Combined loss of *cep290* and *ahi1* does not exacerbate cone degeneration or rhodopsin trafficking defects. Panels show immunohistochemical analysis of dorsal retinas from wild-type (top), *cep290*^*fh297/fh297*^ (middle), and *cep290*^*fh297/fh297*^;*ahi1*^*+/-*^ mutants (bottom) at 6 months of age stained with (A) PNA (red) and Zpr-1 (green) to label cone photoreceptor; (B) Zpr-3 to label rhodopsin; or (C) GRK1 to label rhodopsin kinase. ROS, rod outer segments; COS, cone outer segments; ONL, outer nuclear layer; OPL, outer plexiform layer; INL, inner nuclear layer; IPL, inner plexiform layer; RGC, retinal ganglion cells. Scale bar: 50 μm. (D) Quantification of cone outer segment density or (E) rhodopsin mislocalization from the indicated genotypes at 6 months of age. See methods section for details on quantification. Removing one allele of *ahi1* from a *cep290*^*fh297/fh297*^ mutant background had no effect on cone degeneration or rhodopsin mislocalization. At least 5 unique fish over at least 2 independent experiments were evaluated. \*\**P* < 0.01; *** *P* < 0.0005; **** *P* < 0.0001 as determined by a 1-way ANOVA with a Multiple Comparisons test and Tukey corrections. Data represented as means ± s.e.m.

**Figure 12.**
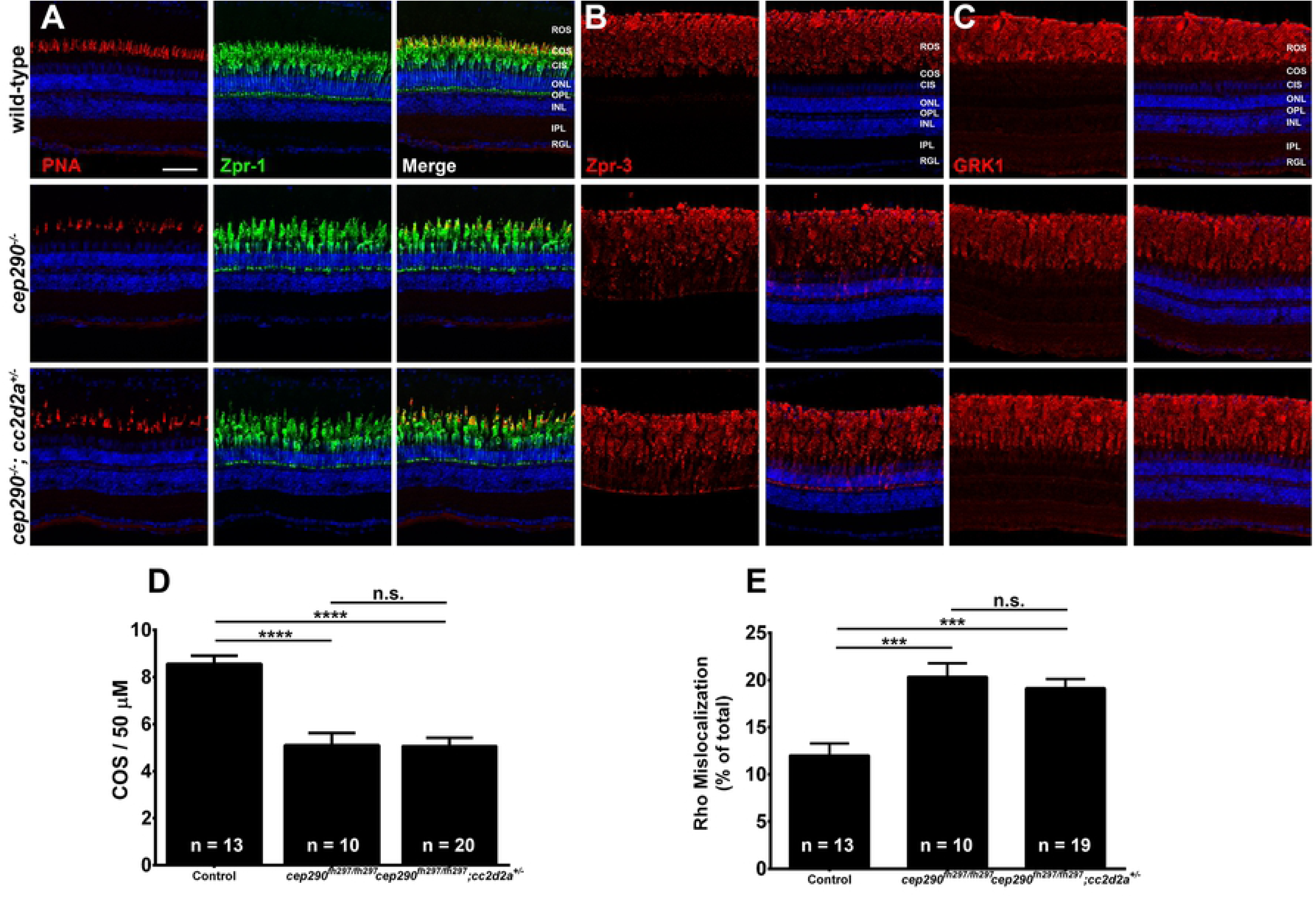
Combined loss of *cep290* and *cc2d2a* does not exacerbate cone degeneration or rhodopsin trafficking defects. Panels show immunohistochemical analysis of dorsal retinas from wild-type (top), *cep290*^*fh297/fh297*^ (middle), and *cep290*^*fh297/fh297*^;*cc2d2a*^*+/-*^ mutants (bottom) at 6 months of age stained with (A) PNA (red) and Zpr-1 (green) to label cone photoreceptor; (B) Zpr-3 to label rhodopsin; or (C) GRK1 to label rhodopsin kinase. ROS, rod outer segments; COS, cone outer segments; ONL, outer nuclear layer; OPL, outer plexiform layer; INL, inner nuclear layer; IPL, inner plexiform layer; RGC, retinal ganglion cells. Scale bar: 50 μm. (D) Quantification of cone outer segment density or (E) rhodopsin mislocalization from the indicated genotypes at 6 months of age. See methods section for details on quantification. Removing one allele of *cc2d2a* from a *cep290*^*fh297/fh297*^ mutant background had no effect on cone degeneration or rhodopsin mislocalization. At least 10 unique fish over at least 2 independent experiments were evaluated. *** *P* < 0.0005; **** *P* < 0.0001 as determined by a 1-way ANOVA with a Multiple Comparisons test and Tukey corrections. Data represented as means ± s.e.m.

**Figure 13.**
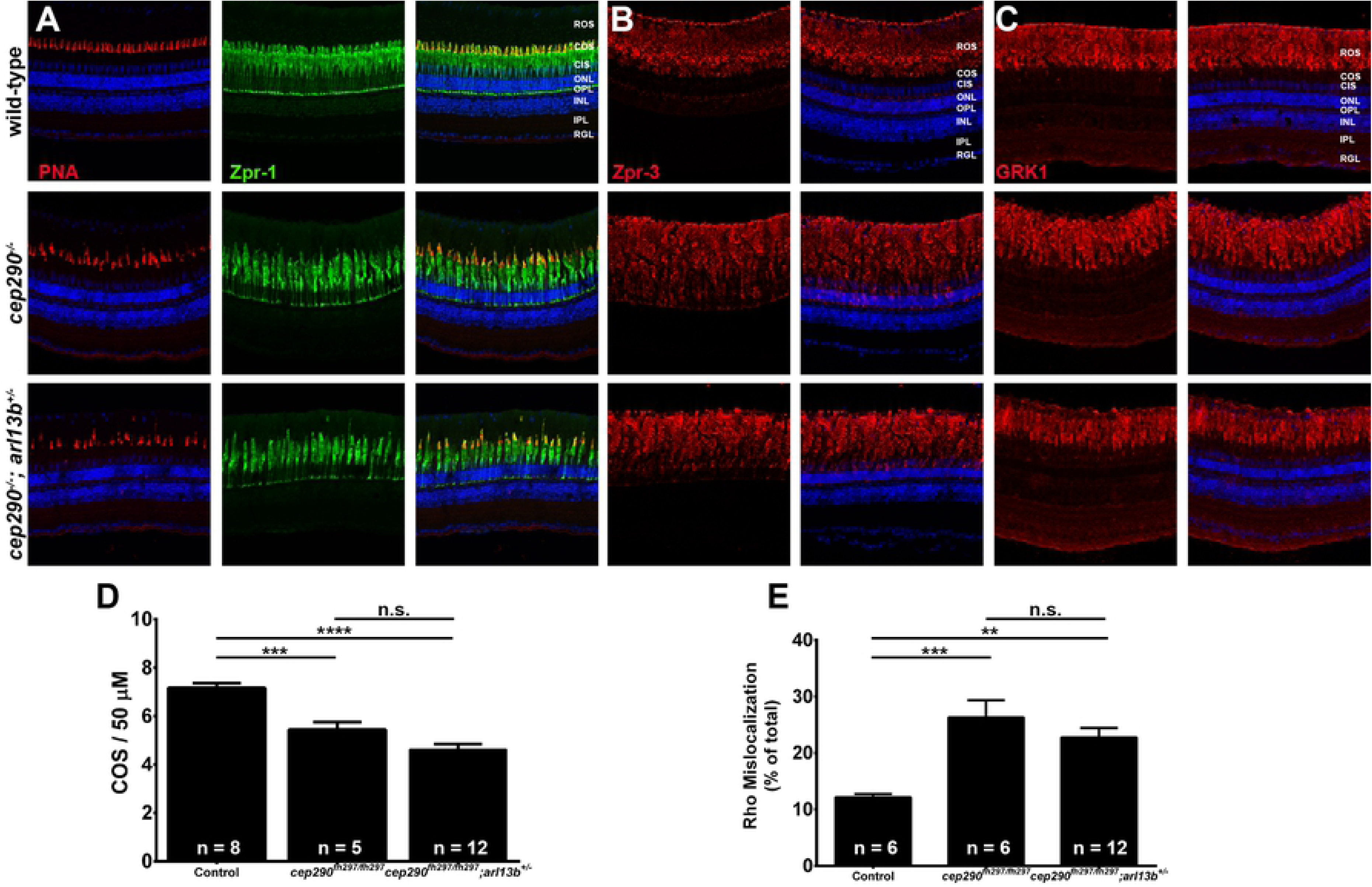
Combined loss of *cep290* and *arl13b* does not exacerbate cone degeneration or rhodopsin trafficking defects. Panels show immunohistochemical analysis of dorsal retinas from wild-type (top), *cep290*^*fh297/fh297*^ (middle), and *cep290*^*fh297/fh297*^;*arl13b*^*+/-*^ mutants (bottom) at 6 months of age stained with (A) PNA (red) and Zpr-1 (green) to label cone photoreceptor; (B) Zpr-3 to label rhodopsin; or (C) GRK1 to label rhodopsin kinase. ROS, rod outer segments; COS, cone outer segments; ONL, outer nuclear layer; OPL, outer plexiform layer; INL, inner nuclear layer; IPL, inner plexiform layer; RGC, retinal ganglion cells. Scale bar: 50 μm. (D) Quantification of cone outer segment density or (E) rhodopsin mislocalization from the indicated genotypes at 6 months of age. See methods section for details on quantification. Removing one allele of *arl13b* from a *cep290*^*fh297/fh297*^ mutant background had no effect on cone degeneration or rhodopsin mislocalization. At least 5 unique fish over at least 2 independent experiments were evaluated. ** *P* < 0.001; *** *P* < 0.0005; **** *P* < 0.0001 as determined by a 1-way ANOVA with a Multiple Comparisons test and Tukey corrections. Data represented as means ± s.e.m.

Finally, we asked if visual function of *cep290*^*fh297/fh297*^ mutant larvae could be diminished by the additional loss of one allele of *ahi1, cc2d2a*, or *arl13b*. We performed pairwise crosses of *cep290*^*+/fh297*^;*ahi1*^*+/-*^, *cep290*^*+/fh297*^;*cc2d2a*^*+/-*^, or *cep290*^*+/fh297*^;*arl13b*^*+/-*^ adults and measured the OKR gain for both contrast sensitivity and visual acuity for all offspring at 5 dpf and subsequently determine the genotype for each animal (Fig. 14). We previously reported that *ahi1*^*-/-*^ mutants have disrupted photoreceptor outer segments but do exhibit normal visual behavior [40]. Although the *cep290*^*fh297/fh297*^;*ahi1*^*+/-*^ mutants had no measurable defect in contrast sensitivity responses (Fig. 14A), a significant reduction in spatial resolution discrimination was observed (Fig. 14B). Interestingly, the *cep290*^*fh297/fh297*^;*cc2d2a*^*+/-*^ mutants had no measurable defect in either contrast sensitivity or spatial resolution responses, although the *cep290*^*fh297/fh297*^;*cc2d2a*^*-/-*^ mutants were more significantly affected (Figs. 14C and D). The *arl13b*^*-/-*^ single mutants showed significant impairment of both contrast sensitivity and spatial frequency discrimination (Figs. 14 E, F). Although the *cep290*^*fh297/fh297*^;*arl13b*^*+/-*^ mutants were not statistically different from *cep290*^*fh297/fh297*^ single mutants in contrast sensitivity, there was a statistically significant difference between *cep290*^*fh297/fh297*^ single mutants and *cep290*^*fh297/fh297*^;*arl13b*^*+/-*^ mutants in spatial resolution (Fig. 14F, purple bar). Interestingly, removing one allele of *cep290* significantly enhanced the defects in both contrast sensitivity and spatial frequency of *arl13b*^*-/-*^ mutants (Figs. 14E, F; blue bars). Taken together, these results suggest that loss of *cep290* is differentially sensitive to the loss of one allele of *ahi1, arl13b,* and *cc2d2a*. The data also suggest that in zebrafish, *arl13b* may not function as a modifier of *cep290*, but *cep290* may instead function as a modifier of *arl13b* in zebrafish. We did not include results from double homozygous mutants because this may reflect an additive effect from two independent phenotypes rather than a true modifier effect. Furthermore, the likelihood that both genes carry two mutant alleles is a highly unlikely to occur in humans with retina disease.

**Figure 14.**
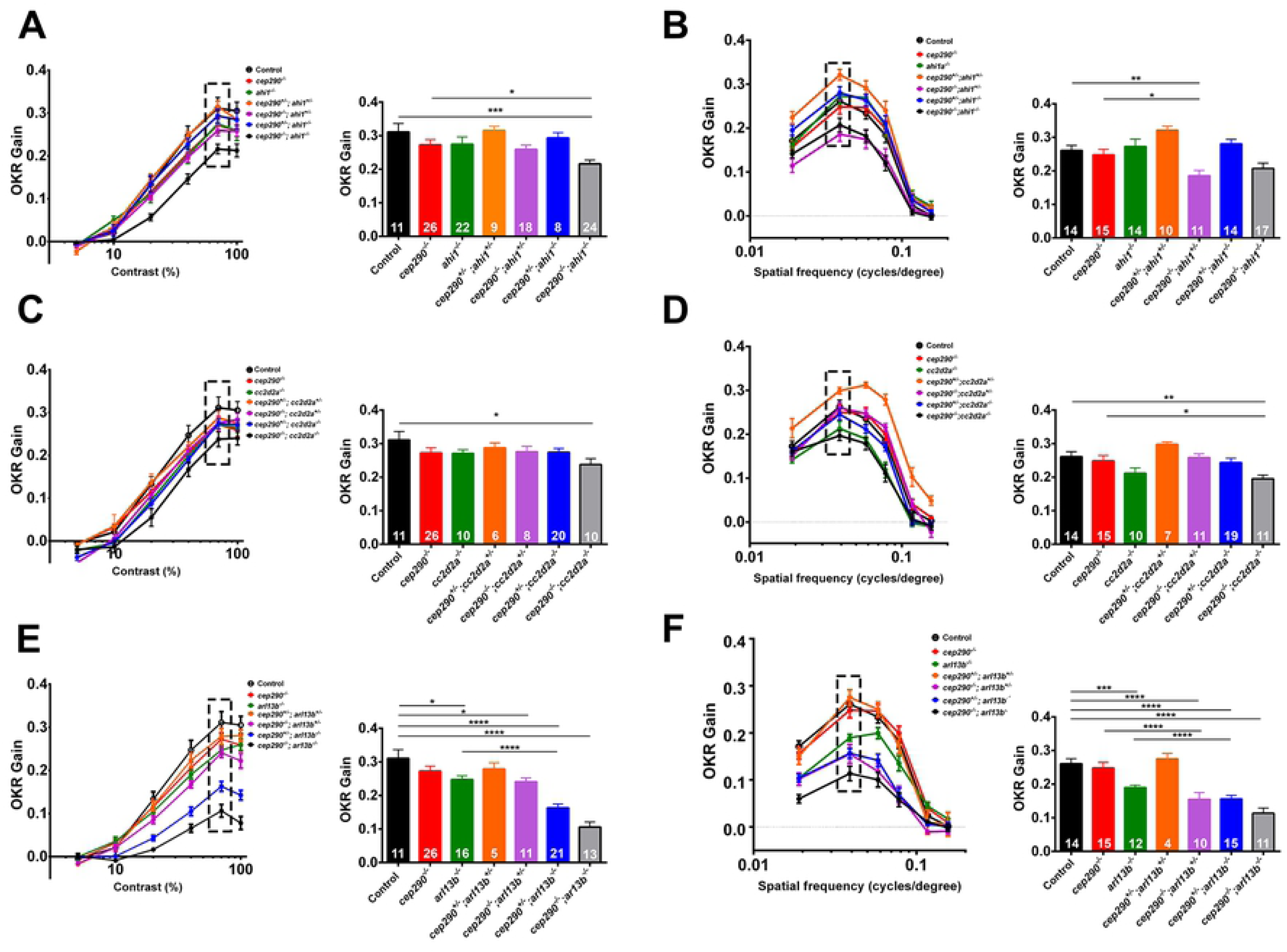
Loss of *ahi1* or *arl13b* impairs visual function of *cep290*^*fh297/fh297*^ mutants. (A, C, E) OKR contrast response function of 5 dpf larvae (left) and bar graph (right) of corresponding data points at the 70% contrast setting (hatched box). (B, D, F) OKR spatial resolution results of 5 dpf larvae (left) and bar graph of corresponding data points at the 0.039 spatial frequency (hatched box). Genotypes are indicated in the legend and in the X-axes. Inset values indicate the total number larvae tested for each genotype. * *P* < 0.05; ** *P* < 0.01; *** *P* < 0.001; \*\*\*\**P* < 0.0001 as determined by a 2-way ANOVA with a Multiple Comparisons test and Tukey corrections. Data are presented as means ± s.e.m.

## DISCUSSION

Mutation of *CEP290* is a major cause of ciliopathies and non-syndromic retinal degeneration. *CEP290* mutations result in a variety of disorders with overlapping but clinically distinct phenotypes and significant phenotypic variation exists between patients diagnosed with the same syndrome. For example, the best corrected visual acuity of several LCA patients with *CEP290* mutations varied from 20/50 to the absence of light perception [9]. The cause of this variation is often attributed to the presence of mutations in second-site modifier genes [33, 71]. Several reports have confirmed a role for modifier genes. For example, an allele of *RPGRIP1L* enhances retinal degeneration across a spectrum of ciliopathies [29] and mutations in *AHI1* were suggested to increase the severity of photoreceptor degeneration in nephronophthisis patients [72]. The effect of genetic modifiers on Cep290 phenotypes has been less clear. In humans, mutations in *AHI1* and *CC2D2A* cause JBTS and both genes have been proposed as potential modifiers of *CEP290* [12, 33]. Missense alleles of *AHI1* were associated with increased neurological involvement in a small number of *CEP290*-LCA patients [33], while morpholino knockdown of *cep290* increased the frequency of kidney cysts in *cc2d2a*^*-/-*^ mutant zebrafish [12]. The absence of one *Bbs4* allele enhanced photoreceptor degeneration in *Cep290*^*rd16/rd16*^ mice [71]. However, mutation of a single allele of *Bbs6* (*Mkks*) rescued the photoreceptor degeneration of *Cep290*^*rd16/rd16*^ mice [73]. Taken together, these data imply that genetic interactions can exert gene- or even allele-specific effects.

In this study, we report that the zebrafish *cep290*^*fh297/fh297*^ mutant retina undergoes progressive cone photoreceptor degeneration beginning at 3 months of age and is accompanied of rhodopsin mislocalization and thinning of the outer nuclear layer by 6 months of age. Retinal development occurs normally and the *cep290*^*fh297/fh297*^ mutants have normal visual acuity as larvae. This is not inconsistent with clinical studies of LCA patients with point mutations in *CEP290*. Younger patients are more likely to have a normal fundus appearance, with older patients showing white flecks or pigmentary retinopathy [8, 33]. These observations indicate that in humans, photoreceptor development is preserved while the long-term photoreceptor survival is affected, similar to what is observed in the *cep290*^*fh297/fh297*^ mutant.

We also investigated how heterozygous mutations in the *ahi1, arl13b,* or *cc2d2a* genes impacted retinal architecture and visual function of *cep290*^*fh297/fh297*^ mutants. Compared to *cep290*^*fh297/fh297*^ mutants, we found that the absence of one allele of these genes did not accelerate retinal degeneration or reduce viability on a *cep290*^*fh297/fh297*^ mutant background. However, loss of one allele of *ahi1* or *arl13b* did decrease the spacial frequency function of *cep290*^*fh297/fh297*^ mutants while the *cep290*^*fh297/fh297*^;*cc2d2a*^*+/-*^ mutants were indistinguishable from *cep290*^*fh297/fh297*^ mutants. We therefore conclude that the retinal phenotypes in zebrafish lacking *cep290* are differentially sensitive to the loss of one allele of *ahi1, arl13b,* or *cc2d2a*. These experiments were prompted by a previous study of zebrafish *cc2d2a*^*-/-*^ mutants that found that injection of morpholinos targeting *cep290* significantly increased the frequency of pronephric cysts at 4 dpf [12], thereby indicating a potential genetic interaction between *cc2d2a* and *cep290* in zebrafish. In a separate report, morpholino-induced knockdown of *cep290* in zebrafish also prevented photoreceptor outer segment formation and other ciliopathy defects [74]. We rarely observed pronephric cysts in *cep290*^*fh297/fh297*^ mutants and the frequency of cysts was not increased by the additional loss of *cc2d2a*, consistent with the lack of genetic interactions in the eye. These contrasting results may reflect the observed differences observed between morpholino-induced phenotypes and mutant phenotypes. Such differences have been attributed to off-target effects of morpholinos or genetic compensation by mutants but not morphants [75, 76].

The slow photoreceptor degeneration observed in the zebrafish *cep290*^*fh297/fh297*^ mutant differs from the phenotypes observed in mice lacking *Cep290*. The *cep290*^*rd16*^ mouse undergoes almost complete loss of rods within 4 weeks of age [20], while a complete knockout of *Cep290* is embryonic lethal [14]. Although the *cep290*^*fh297*^ allele encodes a nonsense mutation, mutant *cep290* transcripts were downregulated by 55% compared to wild-type levels. This is similar to what has been observed in human tissues. Fibroblasts derived from an LCA patient with the c.2991+1665A>G mutation had a 60% reduction in wild-type *CEP290* transcripts that resulted in a corresponding ∼80% reduction in protein levels [43]. A recent study determined that iPSC-derived RPE from a patient with biallelic truncating mutations in *CEP290* maintained protein expression at levels at least 10% of wild-type expression [77]. Furthermore, *CEP290-*LCA patients carrying two nonsense alleles do not undergo the rapid photoreceptor degeneration observed in *Cep290*^*rd16/rd16*^ mice or the increased mortality of the *Cep290*^*ko/ko*^ mice [8, 21]. Several possibilities exist that could explain these differences. It is possible that truncating mutations are subject to exon skipping in humans, thus leading to partial protein production [25]. Exon skipping was not detected in *cep290* transcripts in the zebrafish retina, but perhaps retinal cells differ from other somatic cells in their sensitivity to mutations in the Cep290 gene. We acknowledge that the effect of the *fh297* mutation on protein production in zebrafish remains unknown. Despite multiple attempts with both commercial antibodies [41] and custom-designed antibodies [42], we were unable to detect Cep290 protein in lysates of wild-type or *cep290* mutants by immunoblotting, so the possibility that the mutated gene produces a truncated polypeptide with partial function cannot be excluded. Such partial and truncated polypeptides may also exist in humans with nonsense mutations.

Despite the loss of cone photoreceptors, the rod outer segments appear preserved. Zebrafish typically show a robust capacity to regenerate damaged photoreceptors following acute damage such as intense light exposure, mechanical injury, or chemical-induced toxicity [70, 78-80], but few studies have directly examined whether adult zebrafish have the capacity to regenerate photoreceptors in a model of inherited retinal degeneration. Following retina injury, the Müller glia within the INL undergo a reprogramming event and proliferate as retinal stem cells to regenerate lost neurons. Immunohistochemical analyses with proliferating cell nuclear antigen (PNCA) found a small increase in proliferating cells in the INL of *cep290*^*fh297/fh297*^ mutants, but a significant increase in proliferating cells was seen in the ONL, which have been shown to be rod precursors [69, 81, 82]. It is possible that rods do undergo a slow degeneration in the *cep290*^*fh297/fh297*^ mutants but the dying rods are being continually replaced from the rod progenitor population in the ONL. Because the Müller cells are not proliferative, cone regeneration does not occur. This obviously raises several intriguing questions about how the zebrafish retina differentially responds to acute injury versus a progressive hereditary degeneration.

Zebrafish *cep290* mutants survive to adulthood and reinforce the important role of Cep290 in photoreceptor outer segment maintenance. Furthermore the *cep290*^*fh297/fh297*^ mutant represents a model for slow retinal degeneration that mimics the ocular involvement of *CEP290*- dependent LCA and provides a unique platform to screen for genetic modifiers that accelerate or prevent photoreceptor degeneration. In addition, future work with this model can provide insight into the mechanisms required to trigger photoreceptor regeneration in zebrafish and the signaling pathways required to regenerate lost photoreceptors in *CEP290* patients.

## Acknowledgments

The authors would like to thank Dr. Iain Drummond from Massachusetts General Hospital and members of the Perkins lab for helpful discussions. This work was supported by the National Institutes of Health [NIH R01-EY017037 to B.D.P., NIH P30-EY025585 to Cole Eye Institute]; and the Research to Prevent Blindness [Doris and Jules Stein Professorship to B.D.P.].

## Author Contribution

E. M. Lessieur and B. D. Perkins designed research and analyzed data. E. M. Lessieur, P. Song, G. C. Nivar, J. Fogerty, E. M. Piccillo, R. Rozic and B. D. Perkins performed research. E. M. Lessieur and B. D. Perkins wrote the paper. The authors have no conflicts to declare.

## References

1. Hildebrandt F, Benzing T, Katsanis N. Ciliopathies. N Engl J Med. 2011;364(16):1533-43. Epub 2011/04/22. doi:10.1056/NEJMra1010172. PubMed PMID: 21506742.

2. Sharma N, Berbari NF, Yoder BK. Ciliary dysfunction in developmental abnormalities and diseases. Curr Top Dev Biol. 2008;85:371-427. Epub 2009/01/17. doi:S0070-2153(08)00813-2 [pii] 2. 10.1016/S0070-2153(08)00813-2. PubMed PMID: 19147012.

3. Rachel RA, Li T, Swaroop A. Photoreceptor sensory cilia and ciliopathies: focus on CEP290, RPGR and their interacting proteins. Cilia. 2012;1(1):22. doi:10.1186/2046-2530-1-22. PubMed PMID: 23351659; PubMed Central PMCID: PMCPMC3563624.

4. den Hollander AI, Koenekoop RK, Yzer S, Lopez I, Arends ML, Voesenek KE, et al. Mutations in the CEP290 (NPHP6) gene are a frequent cause of Leber congenital amaurosis. Am J Hum Genet. 2006;79(3):556-61. Epub 2006/08/16. doi:S0002-9297(07)62755-4 [pii] 3. 10.1086/507318. PubMed PMID: 16909394; PubMed Central PMCID: PMC1559533.

5. den Hollander AI, Roepman R, Koenekoop RK, Cremers FP. Leber congenital amaurosis: genes, proteins and disease mechanisms. Prog Retin Eye Res. 2008;27(4):391-419. Epub 2008/07/18. doi:S1350-9462(08)00036-0 [pii] 10.1016/j.preteyeres.2008.05.003. PubMed PMID: 18632300.

6. Coppieters F, Lefever S, Leroy BP, De Baere E. CEP290, a gene with many faces: mutation overview and presentation of CEP290base. Hum Mutat. 2010;31(10):1097–108. doi:10.1002/humu.21337. PubMed PMID: 20690115.

7. Bachmann-Gagescu R, Dempsey JC, Phelps IG, O’Roak BJ, Knutzen DM, Rue TC, et al. Joubert syndrome: a model for untangling recessive disorders with extreme genetic heterogeneity. J Med Genet. 2015;52(8):514–22. doi:10.1136/jmedgenet-2015-103087. PubMed PMID: 26092869.

8. Sheck L, Davies WIL, Moradi P, Robson AG, Kumaran N, Liasis AC, et al. Leber Congenital Amaurosis Associated with Mutations in CEP290, Clinical Phenotype, and Natural History in Preparation for Trials of Novel Therapies. Ophthalmology. 2018. doi:10.1016/j.ophtha.2017.12.013. PubMed PMID: 29398085.

9. Cideciyan AV, Aleman TS, Jacobson SG, Khanna H, Sumaroka A, Aguirre GK, et al. Centrosomal-ciliary gene CEP290/NPHP6 mutations result in blindness with unexpected sparing of photoreceptors and visual brain: implications for therapy of Leber congenital amaurosis. Hum Mutat. 2007;28(11):1074-83. Epub 2007/06/08. doi:10.1002/humu.20565. PubMed PMID: 17554762.

10. Pasadhika S, Fishman GA, Stone EM, Lindeman M, Zelkha R, Lopez I, et al. Differential macular morphology in patients with RPE65-, CEP290-, GUCY2D-, and AIPL1-related Leber congenital amaurosis. Invest Ophthalmol Vis Sci. 2010;51(5):2608–14. doi:10.1167/iovs.09-3734. PubMed PMID: 19959640; PubMed Central PMCID: PMCPMC2868490.

11. Yzer S, Hollander AI, Lopez I, Pott JW, de Faber JT, Cremers FP, et al. Ocular and extra-ocular features of patients with Leber congenital amaurosis and mutations in CEP290. Mol Vis. 2012;18:412-25. PubMed PMID: 22355252; PubMed Central PMCID: PMCPMC3283211.

12. Gorden NT, Arts HH, Parisi MA, Coene KL, Letteboer SJ, van Beersum SE, et al. CC2D2A is mutated in Joubert syndrome and interacts with the ciliopathy-associated basal body protein CEP290. Am J Hum Genet. 2008;83(5):559-71. Epub 2008/10/28. doi:S0002-9297(08)00536-3 [pii] 10.1016/j.ajhg.2008.10.002. PubMed PMID: 18950740; PubMed Central PMCID: PMC2668034.

13. Schouteden C, Serwas D, Palfy M, Dammermann A. The ciliary transition zone functions in cell adhesion but is dispensable for axoneme assembly in C. elegans. J Cell Biol. 2015;210(1):35–44. doi:10.1083/jcb.201501013. PubMed PMID: 26124290; PubMed Central PMCID: PMCPMC4493997.

14. Rachel RA, Yamamoto EA, Dewanjee MK, May-Simera HL, Sergeev YV, Hackett AN, et al. CEP290 alleles in mice disrupt tissue-specific cilia biogenesis and recapitulate features of syndromic ciliopathies. Hum Mol Genet. 2015;24(13):3775–91. doi:10.1093/hmg/ddv123. PubMed PMID: 25859007; PubMed Central PMCID: PMCPMC4459394.

15. Besharse JC, Horst CJ. The photorecetpro connecting cilium. A model for the transition zone. In: Bloodgood RA, editor. Ciliary and Flagellar Membranes. New York: Plenum Publishing Corp.; 1990. p. 389–417.

16. Datta P, Allamargot C, Hudson JS, Andersen EK, Bhattarai S, Drack AV, et al. Accumulation of non-outer segment proteins in the outer segment underlies photoreceptor degeneration in Bardet-Biedl syndrome. Proc Natl Acad Sci U S A. 2015;112(32):E4400–9. doi:10.1073/pnas.1510111112. PubMed PMID: 26216965; PubMed Central PMCID: PMCPMC4538681.

17. Li C, Jensen VL, Park K, Kennedy J, Garcia-Gonzalo FR, Romani M, et al. MKS5 and CEP290 Dependent Assembly Pathway of the Ciliary Transition Zone. PLoS biology. 2016;14(3):e1002416. doi:10.1371/journal.pbio.1002416. PubMed PMID: 26982032; PubMed Central PMCID: PMCPMC4794247.

18. Garcia-Gonzalo FR, Corbit KC, Sirerol-Piquer MS, Ramaswami G, Otto EA, Noriega TR, et al. A transition zone complex regulates mammalian ciliogenesis and ciliary membrane composition. Nat Genet. 2011;43(8):776-84. Epub 2011/07/05. doi:10.1038/ng.8917. ng.891 [pii]. PubMed PMID: 21725307; PubMed Central PMCID: PMC3145011.

19. Murga-Zamalloa CA, Ghosh AK, Patil SB, Reed NA, Chan LS, Davuluri S, et al. Accumulation of the Raf-1 kinase inhibitory protein (Rkip) is associated with Cep290-mediated photoreceptor degeneration in ciliopathies. J Biol Chem. 2011;286(32):28276–86. doi:10.1074/jbc.M111.237560. PubMed PMID: 21685394; PubMed Central PMCID: PMC3151072.

20. Chang B, Khanna H, Hawes N, Jimeno D, He S, Lillo C, et al. In-frame deletion in a novel centrosomal/ciliary protein CEP290/NPHP6 perturbs its interaction with RPGR and results in early-onset retinal degeneration in the rd16 mouse. Hum Mol Genet. 2006;15(11):1847-57. Epub 2006/04/25. doi:ddl107 [pii] 8. 10.1093/hmg/ddl107. PubMed PMID: 16632484.

21. Cideciyan AV, Rachel RA, Aleman TS, Swider M, Schwartz SB, Sumaroka A, et al. Cone photoreceptors are the main targets for gene therapy of NPHP5 (IQCB1) or NPHP6 (CEP290) blindness: generation of an all-cone Nphp6 hypomorph mouse that mimics the human retinal ciliopathy. Hum Mol Genet. 2011;20(7):1411–23. doi:10.1093/hmg/ddr022. PubMed PMID: 21245082; PubMed Central PMCID: PMCPMC3049361.

22. Menotti-Raymond M, David VA, Pflueger S, Roelke ME, Kehler J, O’Brien SJ, et al. Widespread retinal degenerative disease mutation (rdAc) discovered among a large number of popular cat breeds. Vet J. 2010;186(1):32–8. doi:10.1016/j.tvjl.2009.08.010. PubMed PMID: 19747862.

23. Menotti-Raymond M, David VA, Schaffer AA, Stephens R, Wells D, Kumar-Singh R, et al. Mutation in CEP290 discovered for cat model of human retinal degeneration. The Journal of heredity. 2007;98(3):211–20. doi:10.1093/jhered/esm019. PubMed PMID: 17507457.

24. Barny I, Perrault I, Michel C, Soussan M, Goudin N, Rio M, et al. Basal exon skipping and nonsense-associated altered splicing allows bypassing complete CEP290 loss-of-function in individuals with unusually mild retinal disease. Hum Mol Genet. 2018. Epub 2018/05/18. doi:10.1093/hmg/ddy179. PubMed PMID: 29771326.

25. Drivas TG, Wojno AP, Tucker BA, Stone EM, Bennett J. Basal exon skipping and genetic pleiotropy: A predictive model of disease pathogenesis. Sci Transl Med. 2015;7(291):291ra97. doi:10.1126/scitranslmed.aaa5370. PubMed PMID: 26062849; PubMed Central PMCID: PMCPMC4486480.

26. Williams CL, Li C, Kida K, Inglis PN, Mohan S, Semenec L, et al. MKS and NPHP modules cooperate to establish basal body/transition zone membrane associations and ciliary gate function during ciliogenesis. J Cell Biol. 2011;192(6):1023–41. doi:10.1083/jcb.201012116. PubMed PMID: 21422230; PubMed Central PMCID: PMCPMC3063147.

27. Huang L, Szymanska K, Jensen VL, Janecke AR, Innes AM, Davis EE, et al. TMEM237 is mutated in individuals with a Joubert syndrome related disorder and expands the role of the TMEM family at the ciliary transition zone. Am J Hum Genet. 2011;89(6):713–30. doi:10.1016/j.ajhg.2011.11.005. PubMed PMID: 22152675; PubMed Central PMCID: PMC3234373.

28. Yee LE, Garcia-Gonzalo FR, Bowie RV, Li C, Kennedy JK, Ashrafi K, et al. Conserved Genetic Interactions between Ciliopathy Complexes Cooperatively Support Ciliogenesis and Ciliary Signaling. PLoS Genet. 2015;11(11):e1005627. doi:10.1371/journal.pgen.1005627. PubMed PMID: 26540106; PubMed Central PMCID: PMCPMC4635004.

29. Khanna H, Davis EE, Murga-Zamalloa CA, Estrada-Cuzcano A, Lopez I, den Hollander AI, et al. A common allele in RPGRIP1L is a modifier of retinal degeneration in ciliopathies. Nat Genet. 2009. Epub 2009/05/12. doi:ng.366 [pii]9. 10.1038/ng.366. PubMed PMID: 19430481.

30. Sang L, Miller JJ, Corbit KC, Giles RH, Brauer MJ, Otto EA, et al. Mapping the NPHP-JBTS-MKS protein network reveals ciliopathy disease genes and pathways. Cell. 10. 2011;145(4):513–28. doi:10.1016/j.cell.2011.04.019. PubMed PMID: 21565611; PubMed Central PMCID: PMCPMC3383065.

31. Winkelbauer ME, Schafer JC, Haycraft CJ, Swoboda P, Yoder BK. The C. elegans homologs of nephrocystin-1 and nephrocystin-4 are cilia transition zone proteins involved in chemosensory perception. J Cell Sci. 2005;118(Pt 23):5575-87. PubMed PMID: 16291722.

32. Jensen VL, Li C, Bowie RV, Clarke L, Mohan S, Blacque OE, et al. Formation of the transition zone by Mks5/Rpgrip1L establishes a ciliary zone of exclusion (CIZE) that compartmentalises ciliary signalling proteins and controls PIP2 ciliary abundance. EMBO J. 2015;34(20):2537–56. doi:10.15252/embj.201488044. PubMed PMID: 26392567; PubMed Central PMCID: PMCPMC4609185.

33. Coppieters F, Casteels I, Meire F, De Jaegere S, Hooghe S, van Regemorter N, et al. Genetic screening of LCA in Belgium: predominance of CEP290 and identification of potential modifier alleles in AHI1 of CEP290-related phenotypes. Hum Mutat. 2010. Epub 2010/08/05. doi:10.1002/humu.21336. PubMed PMID: 20683928.

34. Dilan TL, Moye AR, Salido EM, Saravanan T, Saravanan K, Goldberg AFX, et al. ARL13B, a Joubert Syndrome-associated protein, is critical for retinogenesis and elaboration of mouse photoreceptor outer segments. J Neurosci. 2018. Epub 2018/12/24. doi:10.1523/JNEUROSCI.1761-18.2018. PubMed PMID: 30573647.

35. Song P, Dudinsky L, Fogerty J, Gaivin R, Perkins BD. Arl13b Interacts With Vangl2 to Regulate Cilia and Photoreceptor Outer Segment Length in Zebrafish. Invest Ophthalmol Vis Sci. 2016;57(10):4517–26. doi:10.1167/iovs.16-19898. PubMed PMID: 27571019; PubMed Central PMCID: PMCPMC5015978.

36. Larkins CE, Aviles GD, East MP, Kahn RA, Caspary T. Arl13b regulates ciliogenesis and the dynamic localization of Shh signaling proteins. Mol Biol Cell. 2011;22(23):4694–703. doi:10.1091/mbc.E10-12-0994. PubMed PMID: 21976698; PubMed Central PMCID: PMC3226485.

37. Littink KW, Pott JW, Collin RW, Kroes HY, Verheij JB, Blokland EA, et al. A novel nonsense mutation in CEP290 induces exon skipping and leads to a relatively mild retinal phenotype. Invest Ophthalmol Vis Sci. 51(7):3646-52. Epub 2010/02/05. doi:iovs.09-5074 [pii] 11. 10.1167/iovs.09-5074. PubMed PMID: 20130272.

38. Daniele LL, Emran F, Lobo GP, Gaivin RJ, Perkins BD. Mutation of wrb, a Component of the Guided Entry of Tail-Anchored Protein Pathway, Disrupts Photoreceptor Synapse Structure and Function. Invest Ophthalmol Vis Sci. 2016;57(7):2942–54. doi:10.1167/iovs.15-18996. PubMed PMID: 27273592; PubMed Central PMCID: PMCPMC4898200.

39. Rinner O, Rick JM, Neuhauss SC. Contrast sensitivity, spatial and temporal tuning of the larval zebrafish optokinetic response. Invest Ophthalmol Vis Sci. 2005;46(1):137-42. PubMed PMID: 15623766.

40. Lessieur EM, Fogerty J, Gaivin RJ, Song P, Perkins BD. The Ciliopathy Gene ahi1 Is Required for Zebrafish Cone Photoreceptor Outer Segment Morphogenesis and Survival. Invest Ophthalmol Vis Sci. 2017;58(1):448–60. doi:10.1167/iovs.16-20326. PubMed PMID: 28118669; PubMed Central PMCID: PMCPMC5270624.

41. Mahuzier A, Gaude HM, Grampa V, Anselme I, Silbermann F, Leroux-Berger M, et al. Dishevelled stabilization by the ciliopathy protein Rpgrip1l is essential for planar cell polarity. J Cell Biol. 2012;198(5):927–40. doi:10.1083/jcb.201111009. PubMed PMID: 22927466; PubMed Central PMCID: PMCPMC3432770.

42. Bergboer JGM, Wyatt C, Austin-Tse C, Yaksi E, Drummond IA. Assaying sensory ciliopathies using calcium biosensor expression in zebrafish ciliated olfactory neurons. Cilia. 2018;7:2. doi:10.1186/s13630-018-0056-1. PubMed PMID: 29568513; PubMed Central PMCID: PMCPMC5856005.

43. Parfitt DA, Lane A, Ramsden CM, Carr AJ, Munro PM, Jovanovic K, et al. Identification and Correction of Mechanisms Underlying Inherited Blindness in Human iPSC-Derived Optic Cups. Cell Stem Cell. 2016;18(6):769–81. doi:10.1016/j.stem.2016.03.021. PubMed PMID: 27151457; PubMed Central PMCID: PMCPMC4899423.

44. Drivas TG, Holzbaur EL, Bennett J. Disruption of CEP290 microtubule/membrane-binding domains causes retinal degeneration. J Clin Invest. 2013;123(10):4525–39. doi:10.1172/JCI69448. PubMed PMID: 24051377; PubMed Central PMCID: PMC3784542.

45. Barbelanne M, Song J, Ahmadzai M, Tsang WY. Pathogenic NPHP5 mutations impair protein interaction with Cep290, a prerequisite for ciliogenesis. Hum Mol Genet. 2013;22(12):2482–94. doi:10.1093/hmg/ddt100. PubMed PMID: 23446637; PubMed Central PMCID: PMCPMC3797088.

46. Krock BL, Perkins BD. The intraflagellar transport protein IFT57 is required for cilia maintenance and regulates IFT-particle-kinesin-II dissociation in vertebrate photoreceptors. J Cell Sci. 2008;121(Pt 11):1907-15. Epub 2008/05/22. doi:121/11/1907 [pii]13. 10.1242/jcs.029397. PubMed PMID: 18492793.

47. Bachmann-Gagescu R, Phelps IG, Stearns G, Link BA, Brockerhoff SE, Moens CB, et al. The ciliopathy gene cc2d2a controls zebrafish photoreceptor outer segment development through a role in Rab8-dependent vesicle trafficking. Hum Mol Genet. 2011;20(20):4041–55. doi:10.1093/hmg/ddr332. PubMed PMID: 21816947; PubMed Central PMCID: PMC3177654.

48. Grimes DT, Boswell CW, Morante NF, Henkelman RM, Burdine RD, Ciruna B. Zebrafish models of idiopathic scoliosis link cerebrospinal fluid flow defects to spine curvature. Science. 2016;352(6291):1341–4. doi:10.1126/science.aaf6419. PubMed PMID: 27284198.

49. Helou J, Otto EA, Attanasio M, Allen SJ, Parisi MA, Glass I, et al. Mutation analysis of NPHP6/CEP290 in patients with Joubert syndrome and Senior-Loken syndrome. J Med Genet. 2007;44(10):657–63. doi:10.1136/jmg.2007.052027. PubMed PMID: 17617513; PubMed Central PMCID: PMCPMC2597962.

50. Wang J, Chang YF, Hamilton JI, Wilkinson MF. Nonsense-associated altered splicing: a frame-dependent response distinct from nonsense-mediated decay. Mol Cell. 2002;10(4):951-7. PubMed PMID: 12419238.

51. Maximov V, Maximova E, Damjanovic I, Maximov P. Detection and resolution of drifting gratings by motion detectors in the fish retina. J Integr Neurosci. 2013;12(1):117–43. doi:10.1142/S0219635213500015. PubMed PMID: 23621461.

52. Purves D AG, Fitzpatrick D, et al. Neuroscience 2nd edition: Sunderland; 2001.

53. Saszik S, Bilotta J, Givin CM. ERG assessment of zebrafish retinal development. Vis Neurosci. 1999;16(5):881–8.

54. Bilotta J, Saszik S, Sutherland SE. Rod contributions to the electroretinogram of the dark-adapted developing zebrafish. Dev Dyn. 2001;222(4):564–70.

55. Vogalis F, Shiraki T, Kojima D, Wada Y, Nishiwaki Y, Jarvinen JL, et al. Ectopic expression of cone-specific G-protein-coupled receptor kinase GRK7 in zebrafish rods leads to lower photosensitivity and altered responses. J Physiol. 2011;589(Pt 9):2321-48. doi:10.1113/jphysiol.2010.204156. PubMed PMID: 21486791; PubMed Central PMCID: PMCPMC3098706.

56. Neuhauss SC, Biehlmaier O, Seeliger MW, Das T, Kohler K, Harris WA, et al. Genetic disorders of vision revealed by a behavioral screen of 400 essential loci in zebrafish. J Neurosci. 1999;19(19):8603–15.

57. Blanks JC, Johnson LV. Specific binding of peanut lectin to a class of retinal photoreceptor cells. A species comparison. Invest Ophthalmol Vis Sci. 1984;25(5):546-57. PubMed PMID: 6715128.

58. Ile KE, Kassen S, Cao C, Vihtehlic T, Shah SD, Mousley CJ, et al. Zebrafish class 1 phosphatidylinositol transfer proteins: PITPbeta and double cone cell outer segment integrity in retina. Traffic. 2010;11(9):1151-67. Epub 2010/06/16. doi:TRA1085 [pii]15. 10.1111/j.1600-0854.2010.01085.x. PubMed PMID: 20545905; PubMed Central PMCID: PMC2919645.

59. Larison KD, Bremiller R. Early onset of phenotype and cell patterning in the embryonic zebrafish retina. Development. 1990;109(3):567–76.

60. Jensen VL, Leroux MR. Gates for soluble and membrane proteins, and two trafficking systems (IFT and LIFT), establish a dynamic ciliary signaling compartment. Curr Opin Cell Biol. 2017;47:83-91. Epub 2017/04/23. doi:10.1016/j.ceb.2017.03.012. PubMed PMID: 28432921.

61. Zhang H, Li S, Doan T, Rieke F, Detwiler PB, Frederick JM, et al. Deletion of PrBP/delta impedes transport of GRK1 and PDE6 catalytic subunits to photoreceptor outer segments. Proc Natl Acad Sci U S A. 2007;104(21):8857–62. doi:10.1073/pnas.0701681104. PubMed PMID: 17496142; PubMed Central PMCID: PMCPMC1885592.

62. Schwarz N, Novoselova TV, Wait R, Hardcastle AJ, Cheetham ME. The X-linked retinitis pigmentosa protein RP2 facilitates G protein traffic. Hum Mol Genet. 2012;21(4):863–73. doi:10.1093/hmg/ddr520. PubMed PMID: 22072390.

63. Zhang H, Hanke-Gogokhia C, Jiang L, Li X, Wang P, Gerstner CD, et al. Mistrafficking of prenylated proteins causes retinitis pigmentosa 2. FASEB J. 2015;29(3):932–42. doi:10.1096/fj.14-257915. PubMed PMID: 25422369; PubMed Central PMCID: PMC4422365.

64. Liu F, Chen J, Yu S, Raghupathy RK, Liu X, Qin Y, et al. Knockout of RP2 decreases GRK1 and rod transducin subunits and leads to photoreceptor degeneration in zebrafish. Hum Mol Genet. 2015;24(16):4648–59. doi:10.1093/hmg/ddv197. PubMed PMID: 26034134.

65. Wan J, Goldman D. Retina regeneration in zebrafish. Curr Opin Genet Dev. 2016;40:41-66. 7. doi:10.1016/j.gde.2016.05.009. PubMed PMID: 27281280; PubMed Central PMCID: PMCPMC5135611.

66. Gorsuch RA, Hyde DR. Regulation of Muller glial dependent neuronal regeneration in the damaged adult zebrafish retina. Exp Eye Res. 2014;123:131–40. doi:10.1016/j.exer.2013.07.012. PubMed PMID: 23880528; PubMed Central PMCID: PMCPMC3877724.

67. Stenkamp DL. The rod photoreceptor lineage of teleost fish. Prog Retin Eye Res. 2011;30(6):395–404. doi:10.1016/j.preteyeres.2011.06.004. PubMed PMID: 21742053; PubMed Central PMCID: PMCPMC3196835.

68. Raymond PA, Rivlin PK. Germinal cells in the goldfish retina that produce rod photoreceptors. Dev Biol. 1987;122(1):120-38. PubMed PMID: 3596007.

69. Otteson DC, D’Costa AR, Hitchcock PF. Putative stem cells and the lineage of rod photoreceptors in the mature retina of the goldfish. Dev Biol. 2001;232(1):62–76.

70. Goldman D. Muller glial cell reprogramming and retina regeneration. Nature reviews Neuroscience. 2014;15(7):431–42. doi:10.1038/nrn3723. PubMed PMID: 24894585; PubMed Central PMCID: PMC4249724.

71. Zhang Y, Seo S, Bhattarai S, Bugge K, Searby CC, Zhang Q, et al. BBS mutations modify phenotypic expression of CEP290-related ciliopathies. Hum Mol Genet. 2014;23(1):40-72. 51. doi:10.1093/hmg/ddt394. PubMed PMID: 23943788; PubMed Central PMCID: PMC3857943.

72. Louie CM, Caridi G, Lopes VS, Brancati F, Kispert A, Lancaster MA, et al. AHI1 is required for photoreceptor outer segment development and is a modifier for retinal degeneration in nephronophthisis. Nat Genet. 2010;42(2):175–80. doi:10.1038/ng.519. PubMed PMID: 20081859; PubMed Central PMCID: PMCPMC2884967.

73. Rachel RA, May-Simera HL, Veleri S, Gotoh N, Choi BY, Murga-Zamalloa C, et al. Combining Cep290 and Mkks ciliopathy alleles in mice rescues sensory defects and restores ciliogenesis. J Clin Invest. 2012;122(4):1233-45. Epub 2012/03/27. doi:10.1172/JCI6098173. 60981 [pii]. PubMed PMID: 22446187; PubMed Central PMCID: PMC3314468.

74. Murga-Zamalloa CA, Atkins SJ, Peranen J, Swaroop A, Khanna H. Interaction of retinitis pigmentosa GTPase regulator (RPGR) with RAB8A GTPase: implications for cilia dysfunction and photoreceptor degeneration. Hum Mol Genet. 2010;19(18):3591–8. doi:10.1093/hmg/ddq275. PubMed PMID: 20631154; PubMed Central PMCID: PMC2928130.

75. Stainier DYR, Raz E, Lawson ND, Ekker SC, Burdine RD, Eisen JS, et al. Guidelines for morpholino use in zebrafish. PLoS Genet. 2017;13(10):e1007000. doi:10.1371/journal.pgen.1007000. PubMed PMID: 29049395; PubMed Central PMCID: PMCPMC5648102.

76. Rossi A, Kontarakis Z, Gerri C, Nolte H, Holper S, Kruger M, et al. Genetic compensation induced by deleterious mutations but not gene knockdowns. Nature. 2015;524(7564):230–3. doi:10.1038/nature14580. PubMed PMID: 26168398.

77. May-Simera HL, Wan Q, Jha BS, Hartford J, Khristov V, Dejene R, et al. Primary Cilium-Mediated Retinal Pigment Epithelium Maturation Is Disrupted in Ciliopathy Patient Cells. Cell reports. 2018;22(1):189–205. doi:10.1016/j.celrep.2017.12.038. PubMed PMID: 29298421.

78. Vihtelic TS, Hyde DR. Light-induced rod and cone cell death and regeneration in the adult albino zebrafish (Danio rerio) retina. J Neurobiol. 2000;44(3):289–307.

79. Maier W, Wolburg H. Regeneration of the goldfish retina after exposure to different doses of ouabain. Cell Tissue Res. 1979;202(1):99-118. PubMed PMID: 509506.

80. Raymond PA, Reifler MJ, Rivlin PK. Regeneration of goldfish retina: rod precursors are a likely source of regenerated cells. Journal of neurobiology. 1988;19(5):431–63. doi:10.1002/neu.480190504. PubMed PMID: 3392530.

81. Johns PR, Fernald RD. Genesis of rods in teleost fish retina. Nature. 1981;293(5828):141–2.

82. Julian D, Ennis K, Korenbrot JI. Birth and fate of proliferative cells in the inner nuclear layer of the mature fish retina. J Comp Neurol. 1998;394(3):271-82. PubMed PMID: 9579393.

